# Centromeric CENP-A loading requires accurate mitotic timing, which is linked to checkpoint proteins

**DOI:** 10.1101/394981

**Authors:** Anne Laure Pauleau, Andrea Bergner, Janko Kajtez, Sylvia Erhardt

## Abstract

A defining feature of centromeres is the presence of the histone H3 variant CENP-A that replaces H3 in a subset of centromeric nucleosomes. In *Drosophila* cultured cells CENP-A deposition at centromeres takes place during the metaphase stage of the cell cycle and strictly depends on the presence of its specific chaperone CAL1. How CENP-A loading is restricted to mitosis is unknown. We found that overexpression of CAL1 is associated with increased CENP-A levels at centromeres and completely uncouples CENP-A loading from mitosis. Moreover, CENP-A levels inversely correlate with mitosis duration. We found that CAL1 interacts with the spindle assembly checkpoint protein and RZZ complex component Zw10 and thus constitutes the anchor for the recruitment of RZZ. Therefore, CAL1 controls CENP-A incorporation at centromeres both quantitatively and temporally, connecting it to the spindle assembly checkpoint to ensure mitotic fidelity.

## Introduction

The formation of two genetically identical daughter cells with a correct and stable genome is of utmost importance during mitosis. Condensed chromosomes are attached and segregated to the opposing poles of the dividing cell at anaphase by the mitotic spindle. At the interface between the chromosomes and the spindle microtubules lies the kinetochore. This multi-protein complex is formed by the components of the KMN network (formed by the Knl1 complex, the Mis12 complex and the Ndc80 complex) (1). The Ndc80 complex is mainly responsible for connecting microtubules with kinetochores while the Knl1 complex primarily functions to coordinate the Spindle Assembly Checkpoint (SAC) (2). The SAC delays entry into anaphase until all chromosomes are properly attached and aligned at the metaphase plate. The metaphase to anaphase transition is controlled by the activation by Cdc20 of the APC/C, a multisubunit ubiquitin ligase that triggers the degradation of cell cycle regulators by the proteasome (3). The SAC proteins Mad2, BubR1, and Bub3 sequester Cdc20 forming the Mitotic Checkpoint Complex (MCC) thereby preventing the activation of the APC/C. In addition, other proteins have been implicated in SAC activity including Bub1, Mad1, the Mps1 and Aurora B kinases and the RZZ complex (formed by the three proteins Rough Deal (ROD), Zw10 and Zwilch) (4). Finally, the Mis12 complex serves as a hub interacting with all kinetochore complexes as well as with the centromere (2).

The kinetochore assembles on centromere during mitosis, a highly specialized chromatin region that is defined by an enrichment of nucleosomes containing the histone H3 variant CENP-A, also called CID in *Drosophila*. In contrast to canonical histones, CENP-A deposition at centromeres is independent of DNA replication and is temporally restricted to a specific cell cycle stage, which varies between organisms: late telophase/early G1 in mammalian cultured cells (5), G2 in S. pombe and plants (6–8), and mitosis to G1 in *Drosophila* (9–12). CENP-A loading requires the action of licensing factors such as the Mis18 complex that modify centromeric chromatin (13), as well as of its dedicated chaperone: HJURP in humans, Scm3 in fungi and CAL1 in *Drosophila* (14–20). Any deregulation of CENP-A and its loading machinery results in the misincorporation of CENP-A into chromosome arm regions. Misincorporated CENP-A is rapidly degraded (21–23). If, however, CENP-A-containing nucleosomes remain at non-centromeric sites ectopic formation of functional kinetochores can occur that may lead to chromosomes segregation defects and aneuploidy (24–26).

The timing of CENP-A is particularly intriguing in *Drosophila* cultured cells as centromeric CENP-A is replenished during prometaphase-metaphase thus coinciding with kinetochore assembly. In *Drosophila melanogaster,* two other proteins are constitutively present at centromeres and essential for centromere function: the conserved protein CENP-C (27) and CENP-A chaperone CAL1. CENP-C has been shown to act as a linker between CENP-A nucleosomes and the Mis12 complex therefore providing a platform for kinetochore assembly (28). CENP-C is also implicated in CENP-A replenishment at centromeres during mitosis by recruiting CAL1 (18, 20). CAL1 interacts with CENP-A in both pre-nucleosomal and nucleosomal complexes (18) and is necessary for CENP-A protein stabilization via Roadkill-Cullin3-mediated mono-ubiquitination (29). Moreover, CAL1 has been previously shown to be the limiting factor for CENP-A centromeric incorporation in fly embryos (30). However, differences in centromere assembly have been reported between *drosophila* cultured cells and tissues. Firstly, CENP-A loading has been shown to take place during mitotic exit in early embryos (9) and during prometaphase to early G1 in cultured cells (10, 12). Second, CENP-C incorporates concomitantly to CENP-A in embryos (9) while this time window seems to be larger in cultured cells (10). We therefore set out to determine more precisely the function of CAL1 in CENP-A loading regulation in *drosophila* cultured cells.

During the course of these studies we found that overexpression of CAL1 not only increases CENP-A abundance at centromeres, it also uncouples CENP-A loading from mitosis. Strikingly, we discovered a co-dependence of mitotic duration and accurate CENP-A loading that is likely mediated by the interaction of the loading machinery of CENP-A with the Spindle Assembly Checkpoint component Zw10. These data suggest an intricate coordination of the spindle assembly checkpoint, CENP-A loading, and mitotic duration.

## Results

### CAL1 levels regulate the amount of CENP-A at centromere

In order to understand whether levels of CAL1 influence CENP-A loading, we overexpressed CAL1 in S2 cells from a copper-inducible pMT promoter. Induced CAL1 localized to centromeres and the nucleolus in a majority of transfected cells (84 %), similar to endogenous CAL1. Cytoplasmic and/or nucleoplasmic CAL1 were visible in about 16 % of the cells (Fig. 1A). Augmented total CAL1 protein levels (2-fold increase; Fig. 1B) led to increased CAL1 levels at centromeres (Fig. 1C) and caused an increase of both endogenous and overexpressed CENP-A (GFP and SNAP) levels (Fig. 1D). Importantly, however, CENP-A was localizing to centromeres only and we did not detect CENP-A at any other chromatin regions on metaphase chromosomes (Fig. 1E-F). This result indicates that CAL1 overexpression does not lead to misincorporation of CENP-A to non-centromeric sites. In contrast to previous observations in embryos (30), centromeric CENP-A levels were increased by up to 2 fold after 24 hours of CAL1 overexpression (Fig. 1G). Surprisingly, CENP-C levels remained largely unaffected by CAL1 overexpression (Fig. 1F-G). These results show that the abundance of CENP-A at centromeres is regulated by the available amount of CAL1.

**Figure 1:**
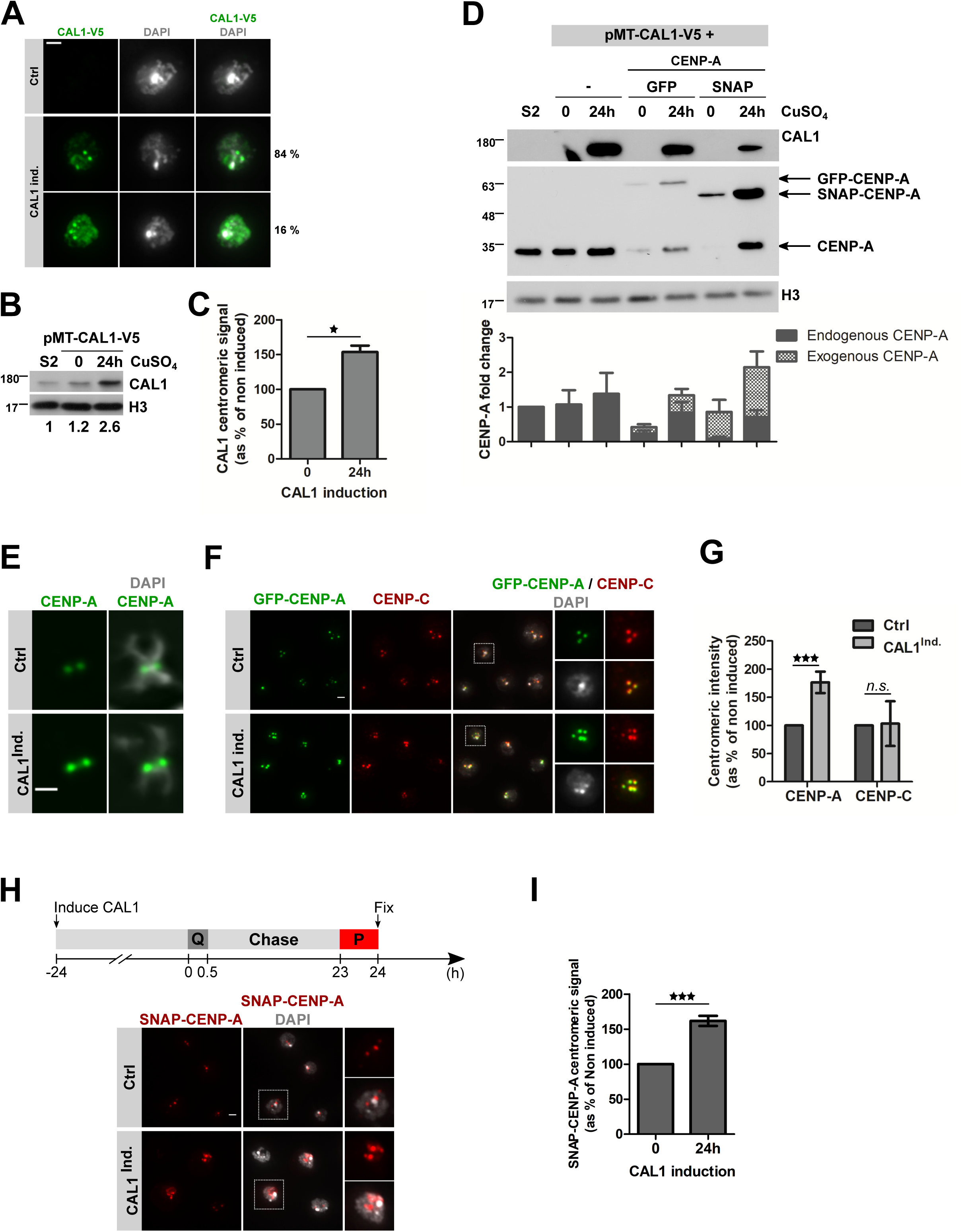
CAL1 overexpression leads to increased CENP-A levels at centromeres. **A.** Immunofluorescence of CAL1-V5 expressing cells. CAL1-V5 expression was induced by addition of 100 µM CuSO_4_ for 24 h. Cells were fixed and stained with anti-V5 antibody (green). DNA was counterstained with DAPI. The percentage of transfected cells displaying either type of localization is shown. **B.** Immunoblot of CAL1-V5 expressing cells treated as in A. Anti-CAL1 (endogenous CAL1 and CAL1-V5) and anti-H3 antibodies were used. **C.** Quantification showing CAL1 centromeric signal intensity per nucleus as % of non-induced control cells. **D.** Immunoblot of CENP-A (either GFP or SNAP N-terminally tagged) expressing cells with or without concomitant CAL1 overexpression. Anti-V5 (CAL1-V5), anti-CENP-A (endogenous CENP-A and tagged-CENP-A) and anti-H3 antibodies were used. The graph shows CENP-A level in each cell line as fold change from baseline levels determined in protein extracts from S2 cells (N=4). **E.** Metaphase chromosomes of CAL1-V5 expressing cells treated as in A. Metaphase chromosomes were stained with anti-CENP-A antibody. DNA was counterstained with DAPI. Scale bar 1 µm. **F.** Immunofluorescence of CAL1-V5/GFP-CENP-A expressing cells treated as in A. Cells were fixed and stained with anti-CENP-C antibody (red). DNA was counterstained with DAPI. **G.** Quantification of F. GFP-CENP-A and CENP-C signal intensities per centromere were measured and are shown as % of non-induced control cells. **H.** Time line of induction experiment followed by a Quench-Chase-Pulse SNAP-tag experiment. After 24 h CAL1-V5 induction, cells were incubated with SNAP-Block to quench existing SNAP-CENP-A molecules (Q), washed and cultured for 24 h. After 24 h chase in presence of CuSO_4_, cells were incubated with SNAP-SiR647 (red) to stain newly synthesized SNAP-CENP-A molecules (P). The images show representative immunofluorescence staining of SNAP-CENP-A expressing cells in control and CAL1 overexpressing cells. DNA was counterstained with DAPI. **I.** Quantification of G. SNAP-CENP-A signal intensity per centromere was measured and is shown as % of non-induced control. All graphs show Mean +/− SEM of 3 experiments (n>300 cells), Student t-test (*n.s*.: non-significant; *: p<0.05; **: p<0.01, ***: p<0.001). Scale bar= 2 µm.

To test whether the increased CENP-A level is caused by increased CENP-A loading we used the SNAP-tag technology in a quench-chase-pulse experiment to label newly synthesized and incorporated CENP-A (5). At day 1 of CAL1 induction, we quenched existing SNAP-CENP-A molecules with a non-fluorescent ligand, cultured the cells for 24 hours to allow approximately one additional cell division, then marked newly synthesized SNAP-CENP-A molecules with a fluorophore and measured fluorescent intensity of newly incorporated SNAP-CENP-A at centromeres (Fig. 1H). Similar to what we observed for GFP-CENP-A, significantly more SNAP-CENP-A incorporated into centromeres in CAL1 overexpressing cells (Fig. 1H-I). These data confirm that CENP-A centromeric levels are regulated by CAL1 and that CAL1 overexpression specifically increases incorporation of newly synthesized CENP-A at centromeric chromatin.

### Overexpression of CAL1 leads to centromeric CENP-A loading outside of mitosis

CENP-A loading takes place during prometaphase-metaphase in *Drosophila* S2 cells (10). To determine if CAL1 overexpression changes CENP-A loading pattern during mitosis, we measured Fluorescence Recovery After Photobleaching (FRAP) of CENP-A, on the assumption that fast recovery of the signal at centromeres corresponds to active loading of CENP-A. We partially bleached GFP-CENP-A signal during prophase and followed recovery of the signal at centromeres until anaphase. We observed a partial recovery of GFP-CENP-A at centromeres in control cells, confirming previous reports that loading of new CENP-A takes place during mitosis (10) (Fig. 2A-B). Surprisingly, GFP-CENP-A recovery rate decreased in CAL1-overexpressing cells compared to control cells (Fig. 2B). One possible explanation for this observation could be that CENP-A centromeric levels get replenished at a different time point in CAL1-overexpressing cells. Indeed, besides loading in mitosis, CENP-A loading has also been reported in G1 in S2 cells (12). We therefore followed cells through mitosis, partially bleached GFP-CENP-A in early G1 and measured GFP-CENP-A signal intensity for 3 hours. However, we did not detect any recovery of the bleached GFP-CENP-A signal (Fig. S1A). We explained these results by the likely absence of any measurable GFP-CENP-A loading in early G1.

**Figure 2:**
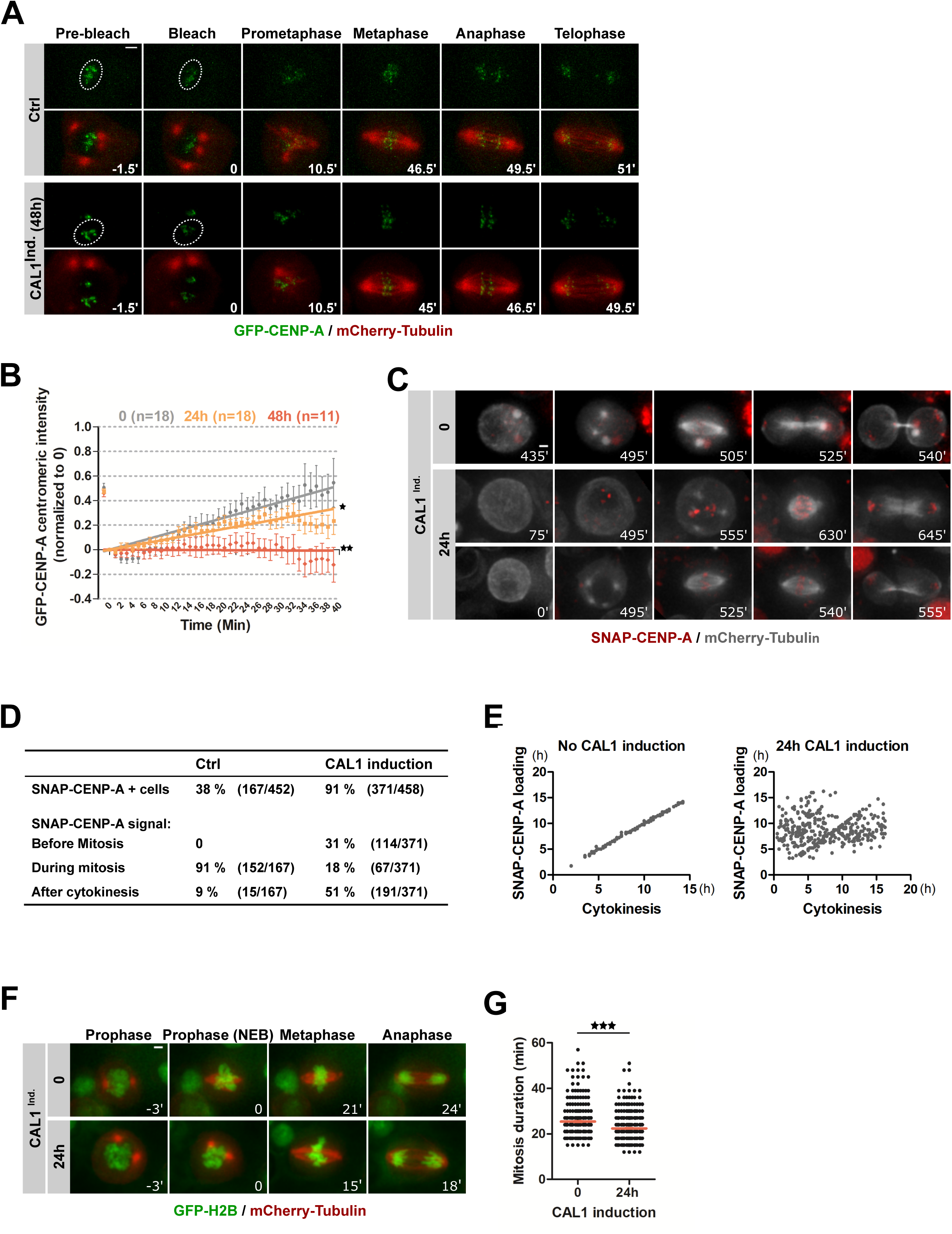
Overexpression of CAL1 leads to CENP-A centromeric loading outside of mitosis. **A.** FRAP experiments of GFP-CENP-A in mitosis in CAL1-overexpressing cells. CAL1-V5 expression was induced for 24 h. GFP-CENP-A signal was partially bleached in prophase and cells were imaged until telophase. Time lapse: 90 s. Scale bar= 2 µm. **B.** Quantification of A. The total GFP-CENP-A centromeric signal is shown as mean +/− SEM, n≥ 11 cells. **C.** Time-lapse imaging of SNAP-CENP-A/mCherry-Tubulin cells with or without concomitant overexpression of CAL1. CAL1-V5 expression was induced for 24 h. The cells were then incubated with SNAP-Block to quench existing SNAP-CENP-A molecules, washed and resuspended in conditional medium containing 0.5 µM SNAP-640 dye. Cells were imaged for 16 h to visualize newly synthesized SNAP-CENP-A. Time lapse 15 min. Scale bar= 2 µm. Intensity levels have been adjusted separately for each condition. **D-E.** Quantifications of C. **D.** Table showing the % of SNAP-CENP-A positive cells (+ cells) and the timing of loading. **E.** For each cycling cell, the earliest detection time point of SNAP-CENP-A (Y-axis) is plotted versus the time of cytokinesis (X-axis). t0 on both axes corresponds to start of imaging after SNAP-Block. **F.** Time-lapse imaging of H2B-GFP/mCherry Tubulin cells with or without concomitant overexpression of CAL1. CAL1-V5 expression was induced for 24 h. The cells were imaged for 16 h. Time lapse 3 min. Scale bar= 2 µm. **G.** Quantification of F showing the mitosis duration from nuclear envelope breakdown (determined by mCherry-Tubulin nuclear diffusion concomitant to DNA condensation) to anaphase entry. Mean +/− SEM, n>200 cells. Student t-test (*: p<0.05; **: p<0.01, ***: p<0.001).

Since the availability of CAL1 seems to be the crucial determinant for CENP-A incorporation, we tested if CAL1-overexpressing cells incorporate CENP-A into centromeric chromatin outside mitosis. Thus, we arrested cells at the metaphase to anaphase transition using the proteasome inhibitor MG132 treatment for 2 hours, blocked existing SNAP-CENP-A molecules using SNAP-Block and cultured the cells for an additional 4 hours in the presence of MG132 (Fig. S1B) similar to a previously published protocol (10). Whereas 10 % of uninduced pMT-CAL1 cells loaded SNAP-CENP-A, the induction of pMT-CAL1 and resulting overexpression of CAL1 led to an incorporation of newly synthesized SNAP-CENP-A in about 40 % of the cells, pointing to loading outside of mitosis (Fig. S1B). To confirm that overexpressed CAL1 indeed loads CENP-A to centromeres outside of mitosis, we followed newly synthesized SNAP-CENP-A live by time-lapse microscopy. We blocked existing SNAP-CENP-A with SNAP-Block, washed and added SNAP-640 fluorophore (31) to the culture medium in order to stain newly synthesized SNAP-CENP-A molecules. Despite some aggregation of the dye, SNAP-CENP-A loading at centromeres was discernable in both CAL1 overexpressing and control cells (Fig. 2C). We only analyzed cycling cells that eventually entered mitosis during the course of imaging and determined whether and when they incorporated SNAP-CENP-A. We found that 38 % of uninduced control cells had SNAP-CENP-A signals at centromeres by the end of the imaging (Fig. 2C-E; video 1). Of those, 91 % loaded SNAP-CENP-A during mitosis and 9 % just after the cell exited mitosis. In contrast, 81 % of cycling CAL1-overexpressing cells had SNAP-CENP-A signals at centromeres (Fig. 2C-E; video 2), and of those 31 % loaded before mitosis and 51 % after the cells exited mitosis. The specificity of the signal was verified by co-staining with anti-CENP-A antibody immediately after block (0 min), and 20 hours of incubation with SNAP-640 (Fig. S1C). No SNAP-CENP-A was detected directly after SNAP-block treatment, confirming the efficient quenching of old SNAP-CENP-A molecules (Fig. S1C). After 20 hours newly synthesized SNAP-CENP-A perfectly co-localized with total CENP-A signals at centromeres. Interestingly, when we counted the percentage of cells that displayed SNAP-CENP-A at centromeres in these fixed samples we did not observe any differences between control and CAL1-overexpressing cells after 20 h (Fig. S1D). However, SNAP-CENP-A centromeric intensity was four times higher in CAL1-overexpressing cells than in control cells (Fig. S1E). This suggests that low SNAP-CENP-A protein levels were masked in the presence of excess SNAP dye in live cell analysis. Nevertheless, the analysis of SNAP-CENP-A loading in cycling cells revealed that cells overexpressing CAL1 load CENP-A throughout the cell cycle while loading outside mitosis does not occur in control cells (Fig. 2D-E). In conclusion, overexpression of the CENP-A-specific chaperone CAL1 uncouples CENP-A centromeric loading from mitosis and leads to an uncontrolled overload of CENP-A at centromeres. Importantly, these live cell analyses revealed that CAL1-overexpressing cells underwent mitosis faster than control cells (Fig. 2A).

To confirm this observation, we performed time-lapse microscopy experiments in cells overexpressing CAL1 together with H2B-GFP and mCherry-Tubulin. Indeed, we found that mitosis duration, defined as the time between partial nuclear envelop breakdown (NEB) and anaphase entry, was 10 % shorter in CAL1-overexpressing cells as compared to control cells (Fig. 2F-G; videos 3-4). Of note, CAL1 overexpressing cells did neither exhibit more mitotic defects (Fig. S2A) nor did they recruit more endogenous kinetochore proteins (CENP-C, Spc105R, BubR1) during prometaphase (Fig. S2B). CAL1-overexpressing cells also display a functional SAC, since cells treated with the microtubule interfering drug Taxol triggered a cell cycle arrest in prometaphase in control cells as well as in CAL1-overexpressing cells (Fig. S2C-D). We concluded that CAL1-overexpressing cells not only incorporated more CENP-A at centromeres, they also progressed faster through mitosis. We therefore wondered next if CENP-A levels could regulate mitosis duration.

### CENP-A levels correlate with mitosis duration

To test the relationship between centromeric CENP-A abundance and the duration of mitosis we designed a strategy to reduce CENP-A levels without inducing chromosome alignment defects that might arrest mitosis (32). We performed CENP-A RNAi depletion in GFP-CENP-A overexpressing cells, which led to undetectable levels of endogenous CENP-A while small amounts of GFP-CENP-A remained in some cells (Fig. 3A-B) and allowed them to go through mitosis (Fig. 3C). Mitosis duration was extended in these cells with reduced CENP-A in comparison to control cells (Fig. 3C-D; videos 5-6). Similar observations have been reported in heterozygous CENP-A mutant fly embryos (33), supporting the hypothesis that CENP-A levels control mitotic length.

**Figure 3:**
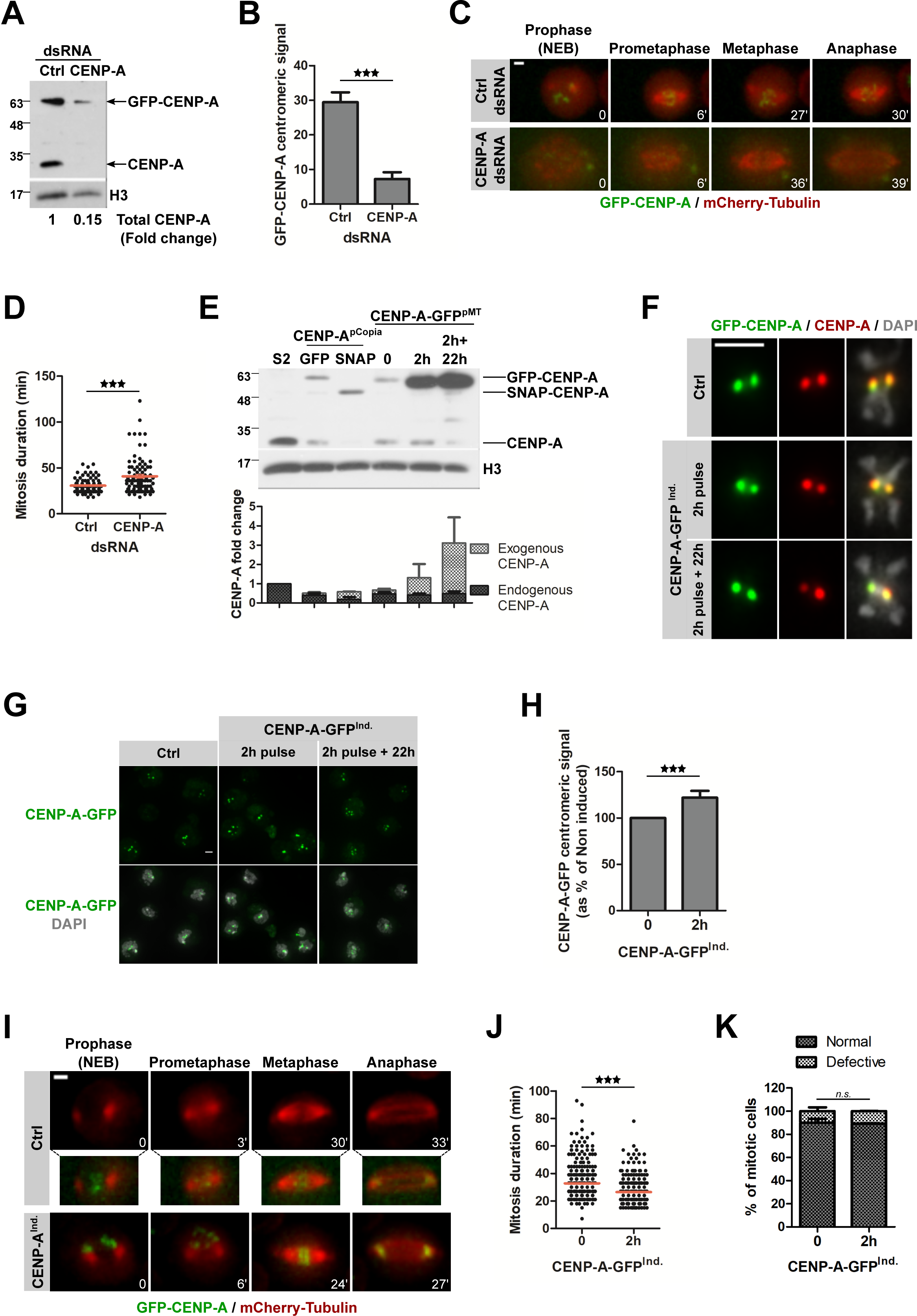
CENP-A levels affects mitosis duration. **A.** Immunoblot showing CENP-A knockdown efficiency in GFP-CENP-A/mCherry-Tubulin expressing cells. An anti-CENP-A antibody was used to detect endogenous CENP-A and overexpressed GFP-CENP-A. **B.** Quantification showing GFP-CENP-A centromeric signal intensity per nucleus at t0 of time-lapse imaging. Mean +/− SEM, n>80 cells. Student t-test (***: p<0.001). Data from 3 experiments were normalized and combined. **C.** Time-lapse imaging of GFP-CENP-A/mCherry-tubulin expressing cells after CENP-A depletion. After 72 h dsRNA treatment, cells were imaged for 16 h. Time lapse 3 min. Scale bar= 2 µm. **D.** Quantification of C. showing the mitosis duration of control and CENP-A depleted cells. Mean +/− SEM, n>80 cells. Student t-test (***: p<0.001). **E.** Immunoblot showing CENP-A levels in the different cell lines used. CENP-A antibody detects endogenous and tagged versions of CENP-A. GFP-CENP-A and SNAP-CENP-A are under the constitutive *copia* promoter. CENP-A-GFP expression was induced by activating the *metallothionein* promoter with 10 µM CuSO_4_ for 2 h. Cells were washed and cultured in conditional medium for 22 h. H3 serves as loading control. The graph shows CENP-A level in each cell line as fold change from baseline levels determined in protein extracts from S2 cells (N=4). **F.** Metaphase chromosomes of CENP-A-GFP expressing cells treated as in E. Metaphase chromosomes were stained with anti-CENP-A antibody. DNA was counterstained with DAPI. Intensities have been adjusted separately for each condition. Scale bar= 2 µm. **G.** Immunofluorescence of CENP-A-GFP cells. CENP-A-GFP expression was induced as in E. DNA was counterstained with DAPI. Scale bar= 2 µm. **H.** Quantification of G. showing the total CENP-A-GFP centromeric intensity per nucleus as % of control. Mean +/− SEM of 3 experiments (n>300 cells), Student t-test (***: p<0.001). **I.** Time lapse imaging of cells expressing mCherry-tubulin and inducible CENP-A-GFP. GFP-CENP-A expression was induced by addition of 10 µM CuSO_4_ to the medium for 2 h. Cells were washed before imaging (16 h). Time lapse 3 min. Scale bar= 2 µm. The intensity levels of CENP-A-GFP in control cells are shown enhanced for visualization purposes. **J.** Quantification of I. showing mitosis duration of control and CENP-A-GFP expressing cells. Mean +/− SEM, n>300 cells. Student t-test (***: p<0.001). **K.** Mitotic phenotypes of CENP-A-GFP/mCherry-Tubulin cells imaged in I. Cells were scored for the accuracy of mitosis: normal or defective (presence of lagging chromosomes in anaphase, formation of tripolar spindles). Mean +/− SEM n > 200 cells. Student t-test (*n.s.*: non-significant).

We next sought to determine if elevated CENP-A levels alone (without CAL1 overexpression) could also influence mitosis duration. However, upregulation of CENP-A to high levels is associated with ectopic CENP-A incorporation into chromosome arms and subsequent defects in mitosis (24). In order to avoid these detrimental effects, we induced CENP-A overexpression from pMT-CENP-A-GFP plasmid with a short and very low CuSO_4_ pulse (2 hours). Under those conditions, CENP-A-GFP was over-expressed while endogenous CENP-A levels were decreased leading to an overall increased CENP-A levels (Fig. 3E). However, 22 hours after the CuSO_4_ pulse untagged CENP-A was almost undetectable, suggesting that cells have a homeostatic mechanism that controls the total abundance of CENP-A protein (Fig. 3E). Under these conditions, non-centromeric CENP-A was undetectable (Fig. 3F) and increased amount of CENP-A-GFP localized to centromeres (Fig. 3G-H). Using time-lapse analysis, we found that cells with increased centromeric CENP-A levels progressed significantly faster through mitosis without increased segregation defects (Fig. 3I-K; videos 7-8). Together these results further suggest that centromeric CENP-A levels influence mitosis duration.

### CENP-A loading correlates with mitosis duration

CENP-A loading peaks during prometaphase-metaphase in *Drosophila* S2 cells. Therefore, we hypothesized that the amount of newly synthesized CENP-A that can be incorporated into centromeric chromatin depends on mitosis duration. Mitosis progression is controlled by the action of SAC proteins, which delay entry into anaphase until all chromosomes are properly attached and aligned at the metaphase plate. We depleted two mitotic checkpoint proteins, BubR1 and Mad2, known to reduce mitosis duration (34–36). These depletions (Fig. S3A) led to a decrease of newly synthesized centromeric SNAP-CENP-A incorporation (Fig. 4A-B). In contrast, depletion of Mis12 (Fig. S3A), which has been previously shown not to affect duration of mitosis in Drosophila cultured cells (37), did not alter SNAP-CENP-A amount at centromeres (Fig. 4A-B). Extending metaphase duration by preventing entry into anaphase through depletion of Spindly (required for silencing of the SAC) (38) or of the APC/C cofactor Cdc27 (39)(Fig. S3A) did not cause an increase of CENP-A levels at centromeres when compared to control (Fig. 4A-B), further confirming that loading occurs prior to anaphase (10). Irreversibly arresting cells in prometaphase by the microtubule-depolymerizing drug Nocodazole or in metaphase by the proteasome inhibitor MG132 also did not cause changes in newly synthesized SNAP-CENP-A levels at centromeres (Fig. S3B-C). SNAP-CAL1 centromeric levels were similarly affected: an acceleration of mitotic timing by BubR1 or Mad2 RNAi led to a decrease of loading, whereas mitotic arrest by Cdc27 or Spindly RNAi did not affect it (Fig. S3D). This set of experiments suggests that the amount of new CENP-A molecules that are incorporated into centromeric chromatin is limited at each round of mitosis and the length of mitosis directly correlates with the amount of centromerically loaded CENP-A (Fig. 4C).

**Figure 4:**
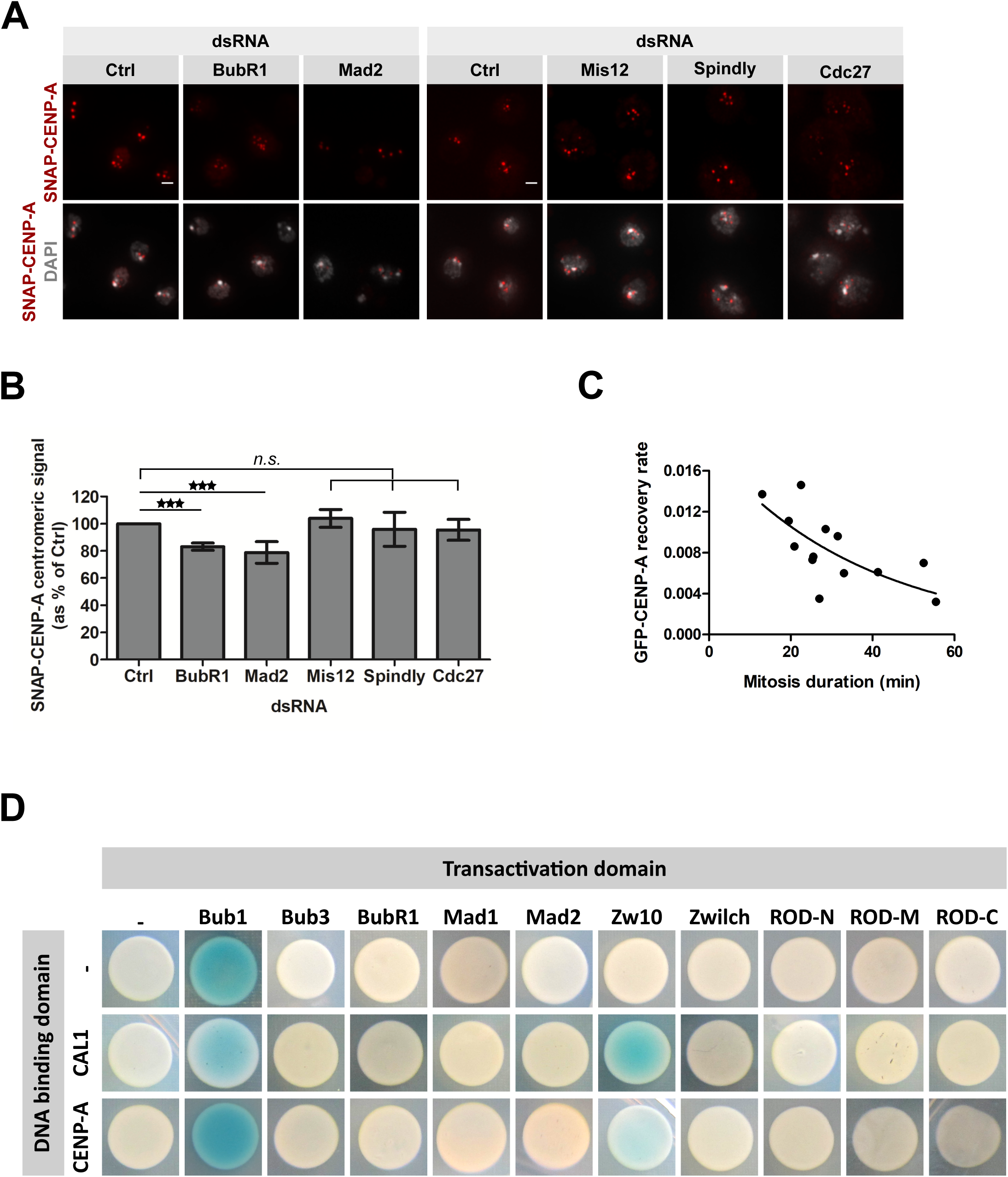
CENP-A loading is correlated with mitosis duration. **A.** SNAP-CENP-A incorporation in BubR1, Mad2, Mis12, Spindly or Cdc27 depleted cells. After 72 h dsRNA treatment, a Quench-Chase-Pulse experiment (scheme in Fig. 1H) was performed to stain newly synthesized SNAP-CENP-A molecules (red). DNA was counterstained with DAPI. Scale bar: 2 µm. **B.** Quantification of A. The total SNAP-CENP-A centromeric signal intensity per nucleus is shown as % of control dsRNA-treated cells. Mean +/− SEM of 3 experiments (n>300 cells), Student t-test (*n.s.*: non-significant; ***: p<0.001). **C.** The recovery rate of GFP-CENP-A after photobleaching in mitosis is plotted against the mitosis duration for each analyzed cell. Pearson’s correlation test, p<0.01. **D.** Yeast two hybrid interaction tests using SAC proteins as prey with either bait CAL1 or CENP-A. Blue color reflects the interaction between the 2 proteins tested.

### RZZ component Zw10 interacts with centromeric proteins

In order to identify a putative connection between CENP-A loading and mitosis progression, we performed a targeted Yeast-two-hybrid screen using either CAL1 or CENP-A as bait and known SAC proteins as prey. These experiments showed that the RZZ component Zw10 weakly but consistently binds to full length CAL1 and CENP-A (Fig. 4D). To identify the domains of each protein required for the interaction, we repeated the YTH analysis with fragments of CAL1 and Zw10. This analysis revealed a strong interaction of the C-terminus of Zw10 (amino acids 482-721) with the C-terminus of CAL1 (amino acids 680-979) (Fig.S4A), the same domain of CAL1 that also binds CENP-C and Roadkill (Fig. S4B) (29, 30). No further interactions were detected between the other checkpoint proteins and the centromeric proteins suggesting that Zw10 might be the key component that connects centromere loading to the checkpoint proteins and ultimately mitosis duration. Our analysis regarding Bub1 has been inconclusive due to self-activation of the Bub1 construct (blue color in the control transformation) but the absence of interaction between CAL1 and Bub1 has been reported previously (30). To validate the interaction between Zw10 and the centromeric proteins, we expressed and purified GST-Zw10, His-CAL1 and His-CENP-A in bacteria. Pulldown experiments using those proteins confirmed that Zw10 directly binds CAL1 (Fig. 5A). We did not detect any interaction between GST-Zw10 and HIS-CENP-A (Fig. S4C). To further confirm the interaction in cells, we established a cell line overexpressing GFP-tagged Zw10. Firstly we confirmed that GFP-Zw10 localization in cultured cells (Fig. S4D) is in accordance with endogenous protein (40–43) as well as transgene expression (44) reported in *Drosophila* tissues: following the disassembly of the nuclear envelope at the onset of mitosis Zw10 was recruited to the kinetochore where it remained until complete chromosome biorientation. GFP-Zw10 was then transported along the spindle microtubules in a process mediated by Spindly and dynein required to silence the SAC (38, 45-47). We next performed immunoprecipitations experiments, which showed that CAL1 co-immunoprecipitated with GFP-Zw10 (Fig. 5B), confirming our pull-down and yeast two hybrid experiments. We concluded from this set of experiments that CAL1 interacts with Zw10, and that this interaction may be involved in coordinating CENP-A centromeric loading with mitosis progression.

**Figure 5:**
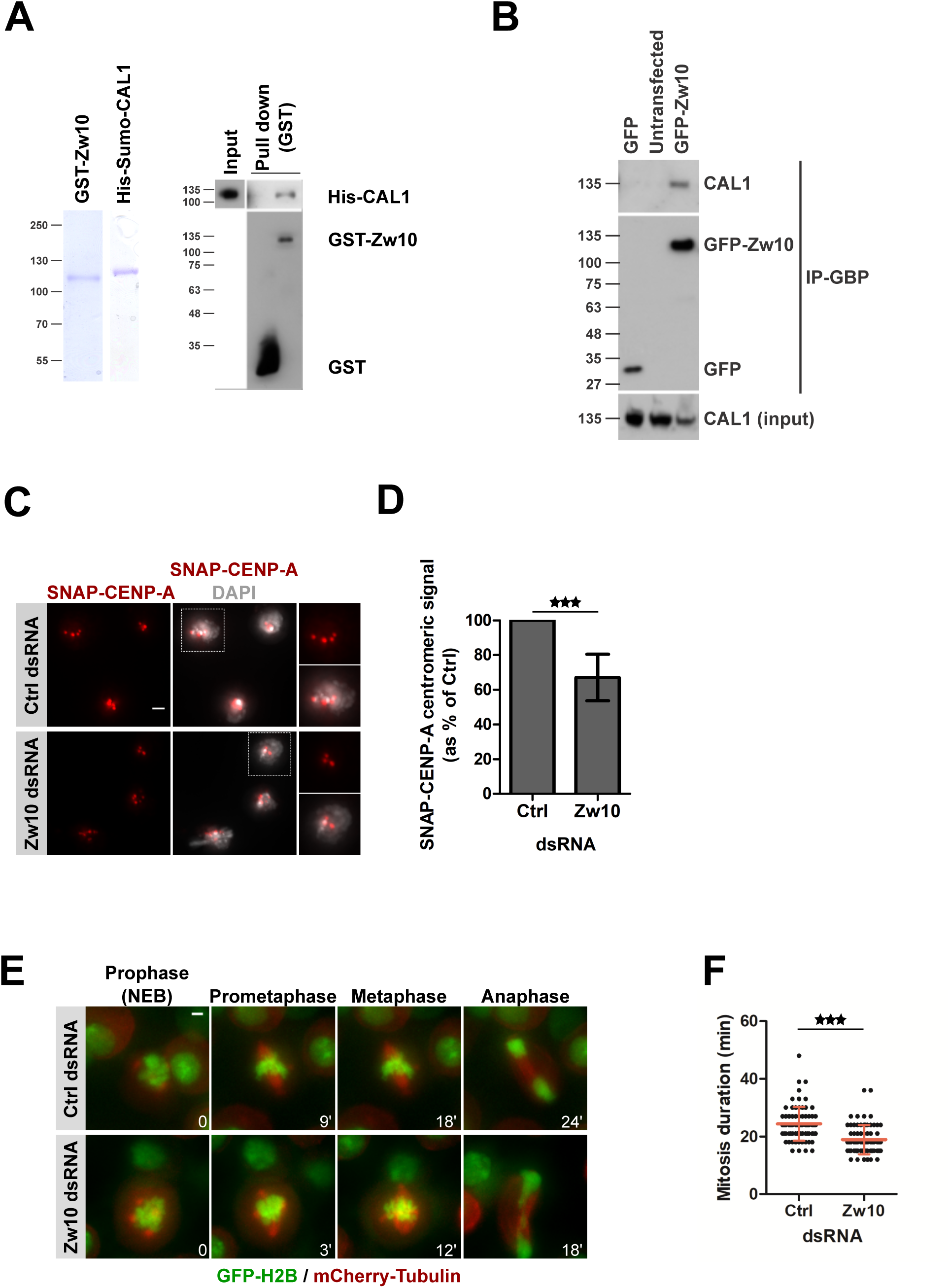
Zw10 interacts with CENP-A loading factor CAL1. **A. *Left panel.*** Coomassie showing purified GST-Zw10 and His-Sumo-CAL1. ***Right panel.*** Immunoblot showing GST-pulldown assays of His-Sumo-CAL1. **B.** Co-immunoprecipitation of CAL1 with GFP-Zw10. GFP (Ctrl) or GFP-Zw10 proteins were immunoprecipitated using the GFP-binding protein (GBP). Pulled down fractions were analyzed by immunoblotting for the presence of CAL1. **C.** SNAP-CENP-A incorporation in Zw10 depleted cells. After 72 h dsRNA treatment, a Quench-Chase-Pulse experiment (scheme in Fig. 1H) was performed to stain newly synthesized SNAP-CENP-A molecules (red). DNA was counterstained with DAPI. Scale bar: 2 µm. **D.** Quantification of C. showing the total SNAP-CENP-A centromeric intensity per nucleus as % of control. Mean +/− SEM of 3 experiments (n>300 cells), Student t-test (***: p<0.001). **E.** Time-lapse imaging of H2B-GFP/mCherry-tubulin expressing cells after Zw10 depletion. After 72 h dsRNA treatment, cells were imaged for 16 h. Time lapse 3 min. Scale bar= 2 µm. **F.** Quantification of G showing the mitosis duration. Mean +/− SEM, n>200 cells. Student t-test (***: p<0.001).

### Zw10 affects the loading of newly synthesized CENP-A during mitosis

The RZZ complex recruits the MAD proteins to the kinetochore, and is thus essential for the SAC (44, 48). Therefore, we sought to determine if Zw10 depletion could recapitulate the effects observed consecutive to the depletion of the other SAC proteins (Fig. 4A-B). Indeed, Zw10 depletion led to a 30 % reduction of newly synthesized SNAP-CENP-A (Fig. 5C-D) and SNAP-CAL1 (Fig. S5A-B) at centromeres compared to control-depleted cells. Time-lapse microscopy analysis of cells stably expressing H2B-GFP and mCherry-Tubulin confirmed that mitotic timing was significantly shorter in Zw10-depleted cells than in control cells (Fig. 5E-F; videos 9-10). This result is in accordance with the Premature Sister Chromatids Separation (PSCS) phenotype observed in Zw10 mutant embryos and larval tissues (41, 49, 50).

To exclude a potential role of Zw10 in CENP-A stability we performed a SNAP-tag pulse chase experiment (Fig. S5C-D) that showed that centromeric SNAP-CENP-A was not affected by Zw10 depletion. Furthermore, FRAP analysis revealed that the recovery rate of GFP-CENP-A at centromeres was comparable in control and Zw10-depleted cells (Fig. S5E-F). Thus, the CENP-A mitotic loading machinery was functional in Zw10-depleted cells suggesting that Zw10 does not influence CENP-A loading directly but through its role in checkpoint activity.

### Zw10 mediates the recruitment of the RZZ to the kinetochore independently of Spc105R

How RZZ is recruited to the kinetochore at mitosis onset remains elusive in *Drosophila*. We first sought to determine if one of the RZZ components is responsible for targeting the whole complex to the kinetochore. We established cell lines overexpressing GFP-ROD and GFP-Zwilch and confirmed that the localization of the tagged proteins is consistent with previous reports (44, 49, 51, 52) (Fig. S4E-F). Knockdown of RZZ complex components (Fig. S6A) led to the expected spindle morphology defects in all cell lines (Fig. 6) (41, 52-54). Importantly, depletion of any complex component led to a reduction of the other GFP-RZZ proteins at kinetochores (Fig. 6) suggesting that the RZZ components are mutually dependent on each other similar to what has been reported in Drosophila tissues (40, 42, 52, 54, 55). However, levels of GFP-Zw10 at kinetochores were the least affected indicating that some GFP-Zw10 is still recruited to kinetochores in the absence of its complex partners ROD or Zwilch (Fig. 6C), suggesting that Zw10 is the most proximal component of the complex at kinetochores in Drosophila cells. Like all kinetochore proteins tested so far (56), the localization of Zw10 to the kinetochore is abolished when CENP-A (Fig. 7B-C) and/or its loading factor CAL1 (Fig. S6B) are depleted, even though Zw10 protein levels remained largely unchanged (Fig. 7A).

**Figure 6:**
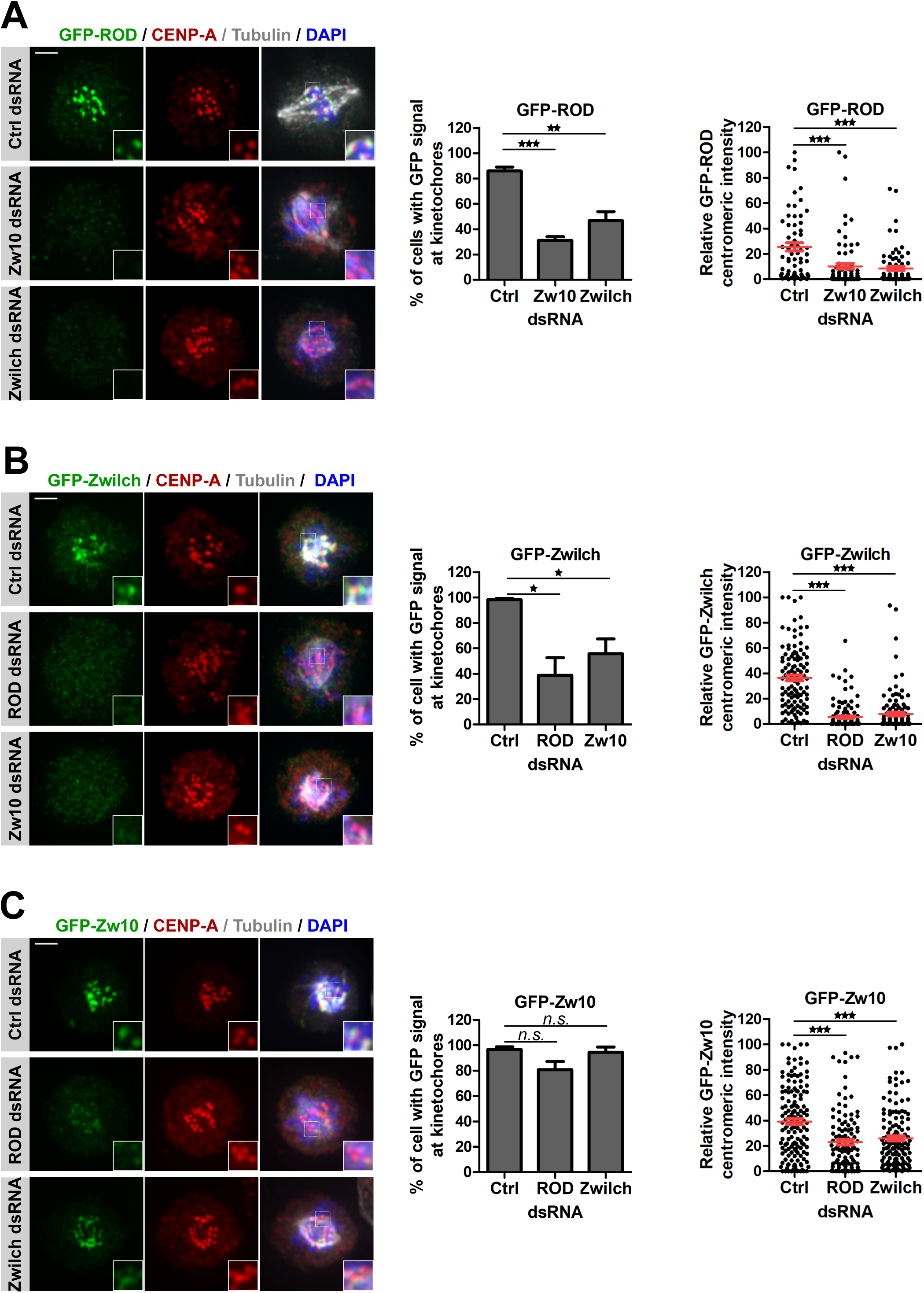
Zw10 is the most proximal RZZ component at kinetochores in *Drosophila*. **A. Left panel** Immunofluorescence of cells expressing GFP-Rod in combination with mCherry-Tubulin after depletion of RZZ components. After 96 h dsRNA treatment, cells were fixed and stained with anti-GFP antibody (green) and anti-CENP-A (red), DNA was counterstained with DAPI. ***Middle panel.*** Prometaphase cells were counted for the presence or absence of GFP signal at kinetochores. ***Right panel.*** Quantification showing the total GFP-Rod kinetochore intensity per cell. GFP fluorescence intensity at kinetochores was measured for each cell and then normalized within one experiment allowing pooling of the measurements for at least 3 experiments per condition. **B.** Similar experiments as in A were performed in GFP-Zwilch overexpressing cells. **C.** Similar experiments as in A were performed in GFP-Zw10 overexpressing cells. Mean +/− SEM, n>100 cells. Student t-test (*n.s.*: non-significant; *: p<0.05; **: p<0.01***: p<0.001). Scale bar= 2 µm.

In mammalian cells, several pathways have been implicated in the recruitment of the RZZ complex including KNL1, Bub1 and Zwint (57–61). As no ortholog of Zwint has been identified in *Drosophila* so far, we tested if the *Drosophila* KNL1 ortholog Spc105R or Bub1 are required for RZZ recruitment to the kinetochore. Depletion of Spc105R (Fig. S6A) did not affect Zw10 localization (Fig. 6C-D), nor did Bub1 depletion (Fig. 6C and S6A, S6C). We also tested Mis12, which has been shown to localize to kinetochores throughout the cell cycle (56) and found that it is also not involved in Zw10 recruitment to the kinetochore (Fig. 6C and S6A, S6D). Since the recruitment of KMN network depends the prior localization of Spc105R and Mis12 at kinetochores (56), those knockdown experiments suggest that the RZZ recruitment to the kinetochore is independent of the KMN network. Conversely, depletion of Zw10 (Fig.7A) did not prevent Spc105R (Fig. 7E-F), Mis12 (Fig. S6E-F), Ndc80 (Fig. 7G-H) or BubR1 (Fig. S6G-H) localization to the kinetochore. Therefore, we propose that the *Drosophila* checkpoint proteins and outer kinetochore components are assembled through two independent branches: on the one hand, CAL1 directly recruits the RZZ complex, which is necessary to localize the MAD proteins to the kinetochore (44, 48), and on the other hand, CENP-C attracts Mis12-Spc105R, which are required to recruit all others components of the KMN network as well as the BUB proteins (28, 56, 62-64). We conclude from these results that there is a direct connection of the SAC and the loading of CENP-A by CAL1, and that the CAL1-RZZ branch directly affects CENP-A loading and mitotic duration, thereby controlling CENP-A loading in a time-dependent manner.

**Figure 7:**
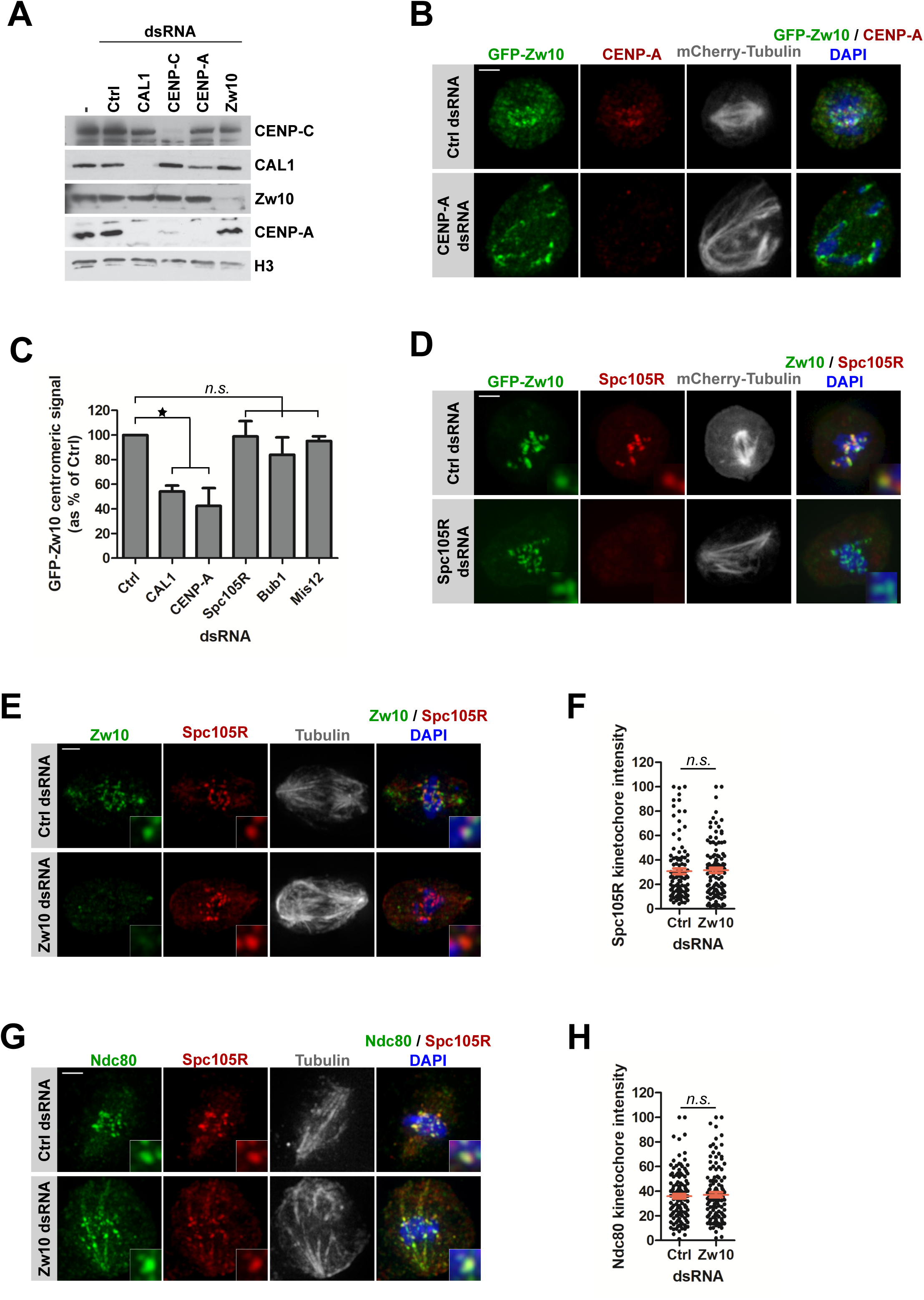
RZZ recruitment to the kinetochore depends on a direct interaction with inner centromere proteins. **A.** Immunoblot showing protein levels after dsRNA treatment. CENP-C, CAL1, Zw10 and CENP-A antibodies were used. H3 serves as a loading control. **B.** Immunofluorescence of GFP-Zw10/mCherry-Tubulin expressing cells after depletion of CENP-A. At day 4 of RNAi, cells were fixed and stained with anti-GFP (green), anti-CENP-A (red) antibodies, DNA was counterstained with DAPI. Scale bar= 2 µm. **C.** Quantifications of B. showing the total GFP-Zw10 centromeric intensity per nucleus as % of control. Prometaphase cells were selected for analysis. Mean +/− SEM of 3 experiments (n>100 cells), Student t-test (*n.s.*: non-significant; *: p<0.05). **D.** Immunofluorescence of GFP-Zw10/mCherry-Tubulin expressing cells after depletion of Spc105R. At day 4 of RNAi, cells were fixed and stained with anti-GFP (green) and anti-Spc105R antibodies (red), DNA was counterstained with DAPI. **E.** Immunofluorescence of S2 cells after depletion of Zw10. At day 4 of RNAi, cells were fixed and stained with anti-Zw10 (green), anti-Spc105R (red) and anti-tubulin (grey) antibodies, DNA was counterstained with DAPI. Scale bar= 2 µm. **F.** Quantification of E. showing the total Spc105R kinetochore intensity per cell. Spc105R fluorescence intensity at kinetochores was measured for each cell and then normalized within one experiment allowing pooling of the measurements for at least 3 experiments per condition. Mean +/− SEM (n>90 cells), Student t-test (n.s.: non-significant) **G.** Immunofluorescence of S2 cells after depletion of Zw10. At day 4 of RNAi, cells were fixed and stained with anti-Ndc80 (green), anti-Spc105R (red) and anti-tubulin (grey) antibodies, DNA was counterstained with DAPI. Scale bar= 2 µm. **H.** Quantification of G. showing the total Ndc80 kinetochore intensity per cell. Ndc80 fluorescence intensity at kinetochores was measured for each cell and then normalized within one experiment allowing pooling of the measurements for at least 3 experiments per condition. Mean +/− SEM (n>90 cells), Student t-test (n.s.: non-significant).

## Discussion

Centromeric chromatin has been shown to behave differently to non-centromeric chromatin in several ways, for instance CENP-A replenishment after DNA synthesis is replication-independent, which depends on specialized loading factors. In *Drosophila* cells, CENP-A loading takes place primarily during prometaphase-metaphase (10). Additional turnover of CENP-A in G1 has been reported leading to the hypothesis that CENP-A could be further incorporated at this stage (12). We did not observe any CENP-A turnover during early G1 in our FRAP experiments when we bleached centromeric CENP-A at the end of cytokinesis. However, our FRAP experiments and our live staining of newly synthesized SNAP-CENP-A confirmed the observations reported by Mellone et al., that the majority of CENP-A loading takes place during mitosis in *Drosophila* cultured cells (10).

In flies, CENP-A incorporation is controlled by its chaperone CAL1 (18, 20, 30). We found that overexpression of CAL1 leads to increased CENP-A protein levels. Under our conditions, CAL1-V5 was strongly overexpressed and caused an intense centromeric and nucleolar localization of CAL1 and a centromere-specific overload of CENP-A. However, ectopic incorporation of CENP-A was never observed suggesting that CAL1 loads CENP-A exclusively to centromeres and that ectopic CENP-A incorporation in flies depends on alternative loading mechanisms similar to what has been suggested in human cells (65). Importantly, increased centromeric CENP-A levels following CAL1 overexpression did not cause chromosome segregation defects but correlated with faster mitosis. A similar acceleration of mitotic timing was observed when CENP-A was overexpressed in a controlled manner that did not lead to massive incorporation at ectopic sites, revealing a possible link between CENP-A loading and mitotic timing. Indeed, shortening mitosis duration by depleting Mad2 or BubR1 (66) was associated with decreased CENP-A loading. Elongating the mitotic time window during which CENP-A is usually loaded (Spindly or Cdc27 depletion, or by drug treatment) did not increase the amount of CENP-A incorporated at centromeres, showing that the length of mitosis alone is insufficient to control CENP-A amounts at centromeres but that only a defined amount of CENP-A can be incorporated at each mitosis, which probably correlates with the available levels of CAL1. Indeed, live analysis of CAL1-overexpressing cells allowed us to visualize newly synthesized CENP-A incorporation to centromeres in all stages of the cell cycle. This strongly suggests that CAL1 controls CENP-A incorporation into centromeric chromatin both quantitatively and temporally. How exactly CENP-A levels at centromeres are sensed is unclear but we identified the RZZ-component Zw10 as a new CAL1 interacting partner, which directly connects CENP-A loading to the SAC. Therefore, we suggest the following model: efficient CENP-A loading by CAL1 during mitosis recruits ZW10 up to a threshold, which is sensed by the SAC. Elevated levels of CENP-A or CAL1, therefore, would accelerate RZZ recruitment and shorten mitosis. It has previously been proposed that BubR1 and Mad2 function as mitotic timer (35, 36, 66, 67). Mad2 but not Mad1 has been shown to be a central determinant of the mitotic timing (66, 68), leading to the hypothesis that cytosolic Mad2-Cdc20 complexes form independently of Mad1 and determine the timing of mitosis (68, 69). Low CENP-A levels at centromeres could lead to more cytosolic Mad2 thereby keeping the timer active longer (33). We hypothesized that higher CENP-A levels at centromeres during early mitosis would accelerate the recruitment of RZZ and consequently Mad2 to the kinetochores, therefore, releasing the timer and shortening mitosis duration in cells where kinetochores attach properly to the spindle microtubules. Interestingly, Nocodazole treatment did not affect CENP-A loading confirming previous observations that kinetochores attachment to the microtubule spindle does not play a role in CENP-A loading (10). These results are pointing further to an additional function of the SAC independent of the control of microtubule attachment. Interestingly, recent evidence shows that RZZ together with Spindly plays a central role in kinetochore expansion during early mitosis to form a fibrous corona that then compacts upon microtubule capture (70–72). Whether and -if so-how the kinetochore expansion by RZZ and spindly is involved in CENP-A loading needs to be investigated in the future.

Many essential components of the SAC require outer kinetochore components for their localization to centromeric regions. In yeast and worms, for instance, Mad1/2 recruitment depends on KNL1 (73, 74). In other eukaryotes, Mad1/Mad2 recruitment to the kinetochore also requires the RZZ complex (58, 59, 75). In mammalian cells, RZZ recruitment to the kinetochore depends on Bub1, which itself requires KNL1/Zwint (59, 61, 76) although some data suggest the existence of a KNL1-independent mechanism (59, 75, 77). Several outer components are missing from the *Drosophila* kinetochore (56, 78) and even though Mad1/2 recruitment to the kinetochore depends on the RZZ complex (48), the factors necessary for the localization of the RZZ to kinetochores are unknown. We showed here that RZZ localization to the kinetochores does not require KNL1^Spc105R^ but depends on the centromeric proteins CAL1 and CENP-A. Therefore, we propose that the *Drosophila* outer kinetochore and components of the SAC assemble through two independent pathways: the CENP-C-KMN-Bub1-Bub3/BubR1 branch (28, 62, 64) or the CAL1-RZZ-Mad1/2 branch. How those two pathways communicate for the formation of MCC complexes remains to be determined. One link may be the KMN complex, since Mad2 has been shown to be diminished in the absence of KMN proteins (37). Interestingly, Spc105R mutation does not affect SAC function in fly embryos suggesting that flies rely more on the RZZ-Mad1/2 branch to engage the SAC (62).

CENP-A expression (79) and its stability (23, 29) together with its dependence on the low abundant and highly specific loading factor CAL1 and the here shown connection to mitotic events are likely interconnected cellular surveillance mechanisms to avoid misincorporation of CENP-A and, therefore, securing genome stability. How CAL1 itself is regulated to obtain such specificity is currently unknown. In conclusion, we show that there is a direct crosstalk between the SAC and the maintenance of centromeric chromatin, ensuring mitotic fidelity not only by controlling microtubule attachment but also by regulating accurate composition of centromeres.

## Materials and methods

### Gene constructs

Plasmid encoding CENP-A tagged with a N-terminal EGFP has been described previously (20) as well as those encoding CENP-A and CAL1 with N-terminal SNAP tag (10). Constructs created for this study (GFP-ROD, GFP-Zwilch, GFP-Zw10) were cloned into AscI and PacI sites of the pCopia–localization and purification (LAP) vector with a basal expression Copia promoter and an N-terminal mEGFP tag (20). For overexpression studies, CAL1-V5 and CENP-A-GFP were cloned into pMT vector (Invitrogen). For yeast two hybrid studies, pMM6 plasmids containing full length CAL1, CAL1 truncations and CENP-A have been described in (29). Bub1, Bub3, BubR1, Mad1, Mad2, Zw10 (full length, aa1-240, aa241-481, aa482-721), Zwilch and Rod (full length, aa1-696, aa697-1392, aa1393-2090) were cloned into pMM5 plasmids. For bacterial expression, pCA528 plasmids containing CAL1 and CENP-A have been described in (29). Zw10 was cloned into pGEX5x-1.

### Cell culture

#### Cell maintenance

*Drosophila* S2 cells were maintained under sterile conditions at 25°C in Schneider medium containing 10 % heat-inactivated fetal bovine serum (FBS) and 100 µg/ml Penicillin/Streptomycin. Expression of genes under the metallothionein promoter (pMT vector) was induced by supplementing the medium with CuSO_4_ (CENP-A-GFP: 10 µM for 2 h; CAL1-V5: 100 µM for 24 h).

#### RNAi and Transfection Experiments

Double-stranded RNA (dsRNA) was produced using the MEGAscript (Ambion) kit according to the manufacturer’s protocol. RNAi was carried out as described previously (20). In short, 10^6^ S2 cells were incubated for 1 h with 15 µg dsRNA in serum-free medium. Subsequently, 2 volumes of 15 % serum containing medium were added. Analysis was usually performed 96 h after treatment.

Stable cell lines were generated by co-transfecting the plasmid of interest together with a plasmid carrying the resistance gene for hygromycin with Cellfectin II (Life Technologies), and the selection process was started 2 d after transfection by supplementing the media with 250 μg/ml Hygromycin B.

#### SNAP quench chase pulse experiments

Cells overexpressing either SNAP-tagged CENP-A or CAL1 proteins were treated with dsRNA as described in RNAi section. After 3 days, cells were collected and incubated in fresh SM containing 10 µM SNAP-Cell Block (New England Biolabs S9106), for 30 min at 25°C, 400 rpm. Cells were then pelleted, washed 3 times (1^st^ wash for 30 min, at 25°C, 400 rpm) and plated in conditional medium. After 24 h, cells were collected and incubated with 4 μM SNAP-Cell TMR-Star (New England Biolabs S9105) or SNAP-SiR647 (New England Biolabs S9102) for 15 min at 25°C, 400 rpm, protected from light. The cells were then pelleted, washed 3 times with SM (1^st^ wash for 30 min, at 25°C, 400 rpm), one time in PBS and prepared for imaging. Cells were settled on a glass coverslip for 10 min, fixed with 4 % PFA for 10 min, washed three times in PBS, and counterstained with DAPI before mounting.

#### SNAP pulse-chase

Cells overexpressing either SNAP-tagged CENP-A or CAL1 proteins were treated with dsRNA as described in RNAi section. After 3 days, cells were collected and incubated in fresh medium containing 4 μM SNAP-Cell TMR-Star (New England Biolabs, Inc.) for 15 min at 25°C, 400 rpm, protected from light. Cells were then washed 3 times with SM (1^st^ wash for 30 min, at 25°C, 400 rpm) and plated in conditional medium. After 24 h, cells were collected and prepared for imaging as previously described.

#### SNAP live imaging

To follow incorporation of newly synthesized SNAP-CENP-A, SNAP-CENP-A was blocked as previously described (see section SNAP quench chase pulse). Cells were then resuspended in conditional medium containing 0.5 µM SNAP-640 dye (31) and placed into an Ibidi imaging chamber before imaging.

### Immunofluorescence

IF was essentially performed as described previously (20). Cells were harvested, washed once with PBS and placed to settle on a glass coverslip for 10 min before fixation with 4 % PFA for 10 min. Cells were washed three times in PBS and permeabilized for 5 min with PBS containing with 0.5 % Triton X-100. Unspecific binding was prevented by blocking the cells for 30 min with 1 % BSA in PBS. Primary antibodies diluted in blocking solution were incubated for 30 min at RT. After three washes with PBS, cells were incubated with the corresponding, fluorescently labeled secondary antibodies (diluted in blocking solution) for 30 min at RT. The secondary antibody incubation and all the following steps were performed while protected from light. After three washes in PBS, DNA was stained for 5 min with DAPI (1 μg/ml in PBS). The cells were then washed two more times with PBS before mounting in Aqua/Polymount medium (Polysciences, Inc.) on a glass slide.

### Preparation of mitotic chromosome spreads

Mitotic chromosome spreads were essentially performed as described previously (80). To obtain mitotic chromosomes, 2×10^5^ exponentially growing cells were arrested in mitosis with 2.5 μg/μl Colcemid for 1 h, centrifuged for 3 min at 800 *g*, resuspended in 0.5 ml hypotonic sodium citrate solution (0.5 % Na-citrate in ddH2O), and incubated for 8–10 min. Cells were spun on positively charged slides in a cytocentrifuge (Shandon 4 Cytospin; Thermo Fisher Scientific) at 900 rpm for 10 min. Slides were placed 10 min in KCM buffer (120 mM KCl, 20 mM NaCl, 10 mM Tris-HCl ph7.7, 0.1 % Triton X-100) before 30 min incubation with primary antibodies diluted in TEEN buffer (1 mM Triethanolamine: HCl pH8.5, 0.2 mM EDTA, 25 mM NaCl, 0.1 % TX-100, 0.1 % BSA) at 37°C. After 3 washes in KB buffer (10 mM Tris-HCl pH7.7, 0.15 M NaCl, 0.1 % BSA) at RT, slides were incubated with secondary antibodies diluted in TEEN buffer for 30 min at 37°C. Slides were then washed 3 times in KB before incubation with 1 µg/ml DAPI diluted in PBS for 5 min at RT. Slides were then mounted in Aqua/Polymount medium (Polysciences, Inc.).

### Microscopy and Image analysis

All Images were acquired on a DeltaVision Core system (Applied Precision) equipped with an Olympus UPlanSApo 100X oil immersion objective (n.a. 1.4) at binning 1×1 (mitotic figures) or 2×2 (interphase). If not stated otherwise, images were taken as z stacks of 0.3 µm increments. All images were deconvolved (20 cycles additive with noise filtering) and maximum-projected using the Applied Precisions soft-WoRx 3.7.1 suite. Image quantification was done using ImageJ software. For determining the number and intensities of centromere/kinetochore dots, a set of plugins developed by the Nikon Imaging Centre, University Heidelberg was used. After marking the nuclear boundaries in the DAPI channel and saving them with the ROI tool, the spots in the channel of interest were enhanced using the DoG spot enhancer plugin. The threshold was adjusted using ‘‘Li’’ instead of default settings. The number of spots with their mean intensities relative to the nuclear area was determined using the ROI particle analyzer plugin.

Time-lapse imaging was performed on a DeltaVision Core system (Applied Precision) equipped either with an Olympus 60X/1.42, N Plan Apo oil immersion objective or an Olympus 40X/1.35, UApo/340 oil objective, at binning 2×2 with 3 min (mitosis duration and fidelity measurements) or 15 min (SNAP-CENP-A live imaging) time-lapse series, for 16 h, with z stacks of 1 µm increment.

FRAP was performed on a Zeiss LSM 780 confocal microscope equipped with a 63x/1.4 oil immersion objective. Number of focal planes per time point, as well as time intervals and total duration were adjusted according to experimental needs. The spacing of focal planes was 1 µm in all experiments. Recovery of EGFP-CENP-A signal was analyzed using FRAP analysis plugin developed by ZMBH imaging facility. After marking the boundaries of the cell of interest, the threshold allowing visualization of centromeric signal exclusively was defined on pre-bleach image. Total intensity of the thresholded signal was then measured for each time point as well as the area. Similar measurements were performed on non-bleached cells from the same area to determine the photobleaching level in each time-series. After correction, EGFP-CENP-A signal intensity was plotted against the time, a linear regression curve was fitted.

### Protein purification and GST-pulldown

Zw10 coding sequence was cloned into pGEX5x-1 bacterial expression vector and transformed into BL21DE3 bacteria. Expression was induced by addition of 1 mM IPTG at OD = 0.6. After 16 h at 21°C, bacteria were pelleted and resuspended in lysis buffer (1 % TX100, 1 mM DTT, 1 mg/ml Lysozyme, 2 mg/ml aprotinin, 5 mg/ml leupeptin, 1 mg/ml pepstatin in PBS). After 30 min incubation at 4°C, bacteria were sonicated (10 cycles, 30 seconds pulse-30 seconds pause, 60 % output power). The lysate was then centrifuged at 15000 g for 45 min. The supernatant was then incubated with Protino glutathione-agarose 4B (Machery-Nagel) for 1 h at 4°C. After 6 washes with 1 % TX100 in PBS, GST-Zw10 was eluted with 20 mM reduced glutathione, 50 mM Tris-HCl pH 8, 1 mM DTT. Buffer was then exchanged using Amicon Ultra 50 K with storage buffer (50 mM Tris-HCl pH 8, 1 mM DTT). His-Sumo-CAL1 and His-Sumo-CENP-A purifications have been described in (29).

Pulldown assays were performed by incubating GST or GST-Zw10 coupled beads with 100 ng His-Sumo-CAL1 or His-Sumo-CENP-A in interaction buffer (20 mM Tris pH 8, 150 mM NaCl, 0.5 mM EDTA, 10 % glycerol, 0.1 % NP-40, 1 mM DTT) for 1 h at 4°C. Beads were then washed 6 times in interaction buffer before resuspension in 1 bead volume 2X sample Laemmli buffer and boiling at 95°C for 5 min.

### Immunoprecipitation

GFP-tagged proteins were isolated using a GFP-specific single chain antibody (GBP) coupled to NHS-activated Sepharose (GE) (81). A total of 1.0–2.0 × 10^8^ cells were lysed in two pellet volumes of 20 mM Tris-HCl pH 7.5, 1 % NP-40, 250 mM NaCl, 20 mM NEM, 10 mM NaF, 2 mM PMSF, 2 mg/ml aprotinin, 5 mg/ml leupeptin, 1 mg/ml pepstatin for 15 min at 4 °C. After clearing the lysate by centrifugation, the supernatant was incubated for 2 h at 4 °C with 50 µl pre-equilibated GBP-beads. The beads were washed six times with lysis buffer and eluted with 2x Laemmli buffer 5 min at 95°C.

### WB analysis and quantification

Cell lysates were separated on a 4-15 % SDS poly-acrylamide gel, transferred onto a nitrocellulose membrane for 2 h at 100 V (Tris-Glycin buffer containing 0.1 % SDS). After blocking in 5 % milk in PBST, primary antibodies were incubated O/N at 4°C in the blocking solution. After washing, secondary antibodies coupled to horseradish peroxidase were added for 1 h at RT before ECL detection (Thermo Fisher Scientific). Immunoblots were quantified using ImageJ.

### Antibodies

The following primary antibodies were used: rabbit H3 (Abcam AB 1791), rabbit CAL1 (29), chicken CENP-A (Erhardt lab + 20), rabbit CENP-A (Active Motif 39713), rabbit YFP (from Pr. Bukau), mouse tubulin (Sigma T9026), mouse V5 (Invitrogen V8012), sheep Scp105R (from Dr. Glover), rabbit Zw10 (from Dr. Goldberg), guinea pig CENP-C (20), rabbit phospho-S10 histone H3 (Abcam ab5176), BubR1 (from Dr. Glover), goat GST (GE healthcare 27457701), rabbit His (Abcam ab9108), rabbit Ndc80 (82), rabbit Mis12 (82). Secondary antibodies coupled to Alexa Fluor 488, Alexa Fluor 546 and Alexa Fluor 647 fluorophores (Molecular Probes) were used for IF, and horseradish peroxidase-conjugated secondary antibodies (Sigma) were used for WB analysis.

### Reverse transcription and Quantitative qPCR

RNA was isolated using Trizol(R) according to the manufacturer’s procedures. Whole genome cDNA synthesis was performed using the Quantitect Reverse Transcription kit (Invitrogen), with a combination of oligo (dT) and random hexamer primers in equal proportions. A control reaction with no reverse transcription was always performed in parallel.

qPCR was performed after cDNA synthesis on a LightCycler 480 (Roche) using LightCycler 480 SYBR Green I Master (Roche). All reactions were run in triplicate in a LightCycler 480 multiwell plate. Actin was used as a reference. The level of each targeted gene in the control mock-treated sample was normalized to 1, and compared with the corresponding–depleted samples.

### Yeast two hybrid

Protein-protein interactions were tested using pMM5 and pMM6 fusion constructs and the yeast strain SGY37VIII. The YTH was performed as described previously (83). In short, interactions were judged based on the activity of β-galactosidase that results in the conversion of X-Gal (5-bromo-4-chloro-3-indolyl-b-D-galactosidase) into a blue dye.

## Acknowledgements

We thank B. Bukau, D. Glover, M. Przewloka, M. Goldberg, R. Karess for antibodies and constructs, V. Belov, A. Butkevich and the Facility of synthetic chemistry of the Max Planck Institute for Biophysical Chemistry (MPIBPC) for the gift of the compound 640SiRH-BG. We thank the ZMBH Imaging Facility and A. Jafar Pour for ImageJ FRAP analysis plugins; H. Lorenz for help with imaging, R. Vlijm and the Erhardt laboratory members for discussions; E. Schiebel and S. Corless for critical comments on the manuscript. We acknowledge funding from the Deutsche Forschungsgemeinschaft through the grant EXC81 (CellNetworks), ER576/2-2, SFB1036, the European Research Council through the grant ERC-CoG-682496 (cenRNA) to SE. ALP is an alumna of the CellNetworks postdoc program.

## Competing interests

No competing interests declared

Online supplemental materials include 6 figures and 10 videos. Fig. S1 shows GFP-CENP-A FRAP in early G1 and SNAP-CENP-A centromeric loading outside mitosis. Fig. S2 shows SAC activity and kinetochore components recruitment in CAL1-overexpressing cells. Fig. S3 shows qPCR results for RNAi efficiency in SNAP-CENP-A cells, SNAP-CENP-A and SNAP-CAL1 quench-chase-pulse experiments in different RNAi and drug treatments. Fig. S4 shows yeast two hybrid with CAL1 and Zw10 fragments, pulldown assay between GST-Zw10 and CENP-A and localization of the GFP-tagged RZZ-proteins during cell cycle. Fig. S5 shows SNAP-CAL1 quench-chase-pulse experiment in Zw10-depleted cells, centromeric SNAP-CENP-A stability assay and GFP-CENP-A FRAP during mitosis in Zw10-depleted cells. Fig. S6 shows qPCR results for RNAi efficiency in GFP-Zw10 cells, GFP-Zw10 localization in CAL1, Bub1 and Mis12-depleted cells, Mis12 and BubR1 localization in Zw10-depleted cells.

## Supplemental figures legends

**Figure S1 related to fig. 1-2.**

**A.** FRAP of GFP-CENP-A in G1 phase. Cells expressing GFP-CENP-A and mCherry-Tubulin were followed through mitosis, GFP-CENP-A signal was partially bleached in early G1 and cells were further imaged for > 2 h. Time lapse: 6 min. Scale bar: 2 µm. **B.** Quantification of A. The total GFP-CENP-A centromeric signal of 6 cells is shown as mean +/− SEM. **C.** Quantification of cells showing SNAP-CENP-A signal at centromeres in control and CAL1-overexpressing cells after 4 h MG132 treatment. CAL1-V5 expression was induced for 24 h. The cells were then incubated for 2 h with MG132 to arrest cells in mitosis prior to SNAP-CENP-A block. A 4-h chase was performed in presence of MG132 to allow synthesis and incorporation of new SNAP-CENP-A into centromeres before incubation with SNAP-Si647 and fixation. **C.** Immunofluorescence of SNAP-CENP-A in control and CAL1 overexpressing cells. CAL1-V5 expression was induced for 24 h. The cells were then incubated with SNAP-Block, washed and resuspended in conditional medium containing SNAP-640 dye. After 20 h, the cells were washed 3 times before fixation and immunostaining with an anti-CENP-A antibody (green). Cells were also taken after block, stained with 0.5 µM SNAP-640 dye for 15 min to check the efficiency of the block. DNA was counterstained with DAPI. Scale bar= 2 µm. **D.** Quantification of C. showing the percentage of cells positive for centromeric SNAP-CENP-A staining. **E.** Quantification of B. showing the total SNAP-CENP-A centromeric intensity per nucleus as % of control. All graphs show Mean +/− SEM of 3 experiments (n>300 cells), Student t-test (n.s.: non-significant; *: p<0.05; **: p<0.01).

**Figure S2 related to fig. 2**

**A.** Mitotic phenotypes of H2B-GFP/mCherry-Tubulin expressing cells with or without concomitant overexpression of CAL1. CAL1-V5 expression was induced for 24 h with 100 µM CuSO_4_ then the cells were imaged for 16 h. Cells were scored for the accuracy of mitosis: normal or defective (presence of lagging chromosomes in anaphase, formation of tripolar spindles). Mean +/− SEM n > 200 cells. Student t-test (*n.s*.: non-significant). **B.** Amount of kinetochore proteins recruited during mitosis in presence or absence of CAL1 overexpression. CAL1-V5 expression was induced for 24 h in GFP-CENP-A/mCherry-Tubulin cells. Cells were then fixed and stained with anti-CENP-C (upper panel, red), anti-Spc105R (middle panel, red) or anti-BubR1 antibody (lower panel, red). DNA was counterstained with DAPI. Scale bar= 2 µm. Kinetochore signal intensity of the indicated proteins is shown as % of control. Only prometaphase cells were analyzed. Mean +/− SEM of 3 experiments (n > 300 cells). Student t-test (*n.s*.: non-significant). **C.** Stills from time-lapse imaging experiments showing GFP-CENP-A/mCherry-Tubulin cells with or without CAL1 overexpression imaged immediately after addition of 1 µM Taxol and after 5 h. CAL1-V5 expression was induced for 24 h. Then 1 µM taxol was added and the cells were imaged for 16 h. Scale bar= 2 µm. **D.** Quantification of C. showing the percentage of cells that arrest (arrest) in response to Taxol treatment or arrest and restart (finish).

**Figure S3 related to fig. 4**

**A.** qPCR results showing mRNA levels after indicated knockdown in SNAP-CENP-A cells as percent of control. **B.** Immunofluorescence of SNAP-CENP-A expressing cells after MG132 or Nocodazole treatment. A Quench-Chase-Pulse experiment was performed to stain newly synthesized SNAP-CENP-A molecules. The 24 h chase was performed in presence or absence of the indicated drug. Cells were stained with an anti-S10-phospho-H3 antibody to identify mitotic cells. DNA was counterstained with DAPI. Scale bar= 2 µm. **C.** Quantification of B. showing the SNAP-CENP-A centromeric intensity as % of control. Only Phospho-H3 positive cells were analyzed. Mean+/− SEM of 3 experiments (n>300 cells), Student t-test (n.s= non-significant). **D.** Quantification showing the total SNAP-CAL1 centromeric intensity per nucleus as % of control after knockdown of the indicated proteins. Mean+/− SEM of 3 experiments (n>300 cells), Student t-test (**: p<0.01; n.s.: non-significant).

**Figure S4 related to fig. 5**

**A.** Yeast two hybrid interaction tests. Blue color reflects the interaction between the 2 proteins tested. **B.** Scheme showing CAL1 functional domains and its binding to its known partners. BD: binding domain. **C. *Left panel.*** Coomassie showing purified His-Sumo-CENP-A. ***Right panel.*** Pulldown assay between GST or GST-Zw10 with His-Sumo-CENP-A. **D-F.** Localization of GFP-Zw10 (**D**), GFP-Zwilch (**E**), GFP-ROD (**F**) during cell cycle. Cells expressing each GFP-RZZ component concomitant with mCherry-Tubulin were fixed and stained with anti-GFP (green) and anti-CENP-A (red) antibodies, DNA was counterstained with DAPI. Scale bar= 2 µm.

**Figure S5 related to fig. 5**

**A.** Immunofluorescence of SNAP-CAL1 expressing cells after Zw10 depletion. After 72 h dsRNA treatment, a Quench-Chase-Pulse experiment (scheme in Fig. 1H) was performed to stain newly synthesized SNAP-CAL1 molecules. DNA was counterstained with DAPI. Scale bar= 2 µm. **B.** Quantification of A. showing the total SNAP-CAL1 centromeric intensity per nucleus as % of control. Mean +/− SEM of 3 experiments (n>300 cells), Student t-test (**: p<0.01). **C.** Immunofluorescence of SNAP-CENP-A expressing cells after Zw10 depletion. After 72 h dsRNA treatment, cells were incubated with TMR-Star (P) to stain existing SNAP-CENP-A molecules, washed and put back in culture (t0). After 24 h chase, cells were fixed (t24). Note that no SNAP-Block was performed for this experiment. DNA was counterstained with DAPI. Scale bar= 2 µm. **D.** Quantification of C. showing the total SNAP-CENP-A centromeric intensity per nucleus as % of t0. Mean +/− SEM of 3 experiments (n>300 cells), Student t-test (n.s= non-significant). **E.** FRAP of GFP-CENP-A in mitosis after Zw10 depletion. After 72 h dsRNA treatment, GFP-CENP-A signal was partially (about 50-60 %) bleached in prophase and cells were imaged until telophase. Time lapse: 90 s. Scale bar= 2 µm. **F.** Quantification of E. The total GFP-CENP-A centromeric signal of at least 8 cells is displayed as mean +/− SEM.

**Figure S6 related to fig. 6-7**

**A.** qPCR results showing mRNA levels after indicated knockdown in GFP-Zw10 cells as percent of control. **B-D.** Immunofluorescence of GFP-Zw10/mCherry-Tubulin expressing cells after depletion of CAL1 (**B**), Bub1 (**C**) or Mis12 (**D**). After 96 h dsRNA treatment, cells were fixed and stained with anti-GFP (green) and anti-CENP-A (B) or Scp105R (C-D)(red) antibodies, DNA was counterstained with DAPI. Scale bar= 2 µm. **E.** Immunofluorescence of S2 cells after depletion of Zw10. At day 4 of RNAi, cells were fixed and stained with anti-Mis12 (green), anti-Spc105R (red) and anti-tubulin (grey) antibodies, DNA was counterstained with DAPI. Scale bar= 2 µm. **F.** Quantification of E. showing the total Mis12 kinetochore intensity per mitotic cell. Mis12 fluorescence intensity at kinetochores was measured for each cell and then normalized within one experiment allowing pooling of the measurements for at least 3 experiments per condition. Mean +/− SEM (n>90 cells), Student t-test (n.s.: non-significant). **G.** Immunofluorescence of S2 cells after depletion of Zw10. At day 4 of RNAi, cells were fixed and stained with anti-BubR1 (green) and anti-tubulin (grey) antibodies, DNA was counterstained with DAPI. Scale bar= 2 µm. **H.** Quantification of G. showing the total BubR1 kinetochore intensity per mitotic cell. BubR1 fluorescence intensity at kinetochores was measured for each cell and then normalized within one experiment allowing pooling of the measurements for at least 3 experiments per condition. Mean +/− SEM (n>90 cells), Student t-test (n.s.: non-significant).

## Supplemental Videos

**<u>Video 1 (related to Figure 2D).</u>**

SNAP-CENP-A/mCherry-tubulin expressing cells containing inducible CAL1-V5. No CAL1-V5 induction. SNAP-CENP-A molecules were quenched before addition of 0.5 µM SNAP-640 dye. Imaging was performed for 16 h on a Deltavision microscope (60X, Bin 2), at 25°C. Time lapse 15 min. Scale bar= 5 µm.

**<u>Video 2 (related to Figure 2D):</u>**

SNAP-CENP-A/mCherry-tubulin expressing cells containing inducible CAL1-V5. CAL1-V5 expression was induced for 24 h with 100 µM CuSO_4_. SNAP-CENP-A molecules were quenched before addition of 0.5 µM SNAP-640 dye. Imaging was performed for 16 h on a Deltavision microscope (60X, Bin 2), at 25°C. Time lapse 15 min. Scale bar= 5 µm.

**<u>Video 3 (related to Figure 2G):</u>**

H2B-GFP/mCherry-tubulin expressing cells containing inducible CAL1-V5. No CAL1-V5 induction. Imaging was performed for 16 h on a Deltavision microscope (60X, Bin 2), at 25°C. Time lapse 3 min. Scale bar= 5 µm.

**<u>Video 4 (related to Figure 2G):</u>**

H2B-GFP/mCherry-tubulin expressing cells containing inducible CAL1-V5. CAL1-V5 expression was induced for 24 h prior to imaging with 100 µM CuSO_4_. Imaging was performed for 16 h on a Deltavision microscope (60X, Bin 2), at 25°C. Time lapse 3 min. Scale bar= 5 µm.

**<u>Video 5 (related to Figure 3C):</u>**

GFP-CENP-A/mCherry-Tubulin expressing cells after Ctrl dsRNA treatment (72 h). Imaging was performed for 16 h on a Deltavision microscope (40X, Bin 2), at 25°C. Time lapse 3 min. Scale bar= 5 µm.

**<u>Video 6 (related to Figure 3C):</u>**

GFP-CENP-A/mCherry-Tubulin expressing cells after CENP-A dsRNA treatment (72 h). Imaging was performed for 16 h on a Deltavision microscope (40X, Bin 2), at 25°C. Time lapse 3 min. Scale bar= 5 µm.

**<u>Video 7 (related to Figure 3I):</u>**

CENP-A-GFP/mCherry-tubulin expressing cells. Visible CENP-A-GFP results from leaky metallothionein promoter. Imaging was performed for 16 h on a Deltavision microscope (40X, Bin 2), at 25 °C. Time lapse 3 min. Scale bar= 5 µm.

**<u>Video 8 (related to Figure 3I):</u>**

CENP-A-GFP/mCherry-tubulin expressing cells. CENP-A-GFP expression was induced for 2 h with 10 µM CuSO_4_ then washed 3 times before being placed in Ibidi chamber. Imaging was performed for 16 h on a Deltavision microscope (40X, Bin 2), at 25°C. Time lapse 3 min. Scale bar= 5 µm.

**<u>Video 9 (related to Figure 5E):</u>**

H2B-GFP/mCherry-tubulin expressing cells after Ctrl dsRNA treatment (72 h). Imaging was performed for 16 h on a Deltavision microscope (60X, Bin 2), at 25°C. Time lapse 3 min. Scale bar= 5 µm.

**<u>Video 10 (related Figure 5E):</u>**

H2B-GFP/mCherry-tubulin expressing cells after Zw10 dsRNA treatment (72 h). Imaging was performed for 16 h on a Deltavision microscope (60X, Bin 2), at 25°C. Time lapse 3 min. Scale bar= 5 µm.

