## Supplementary figures and images for "Centromeric CENP-A loading requires accurate mitotic timing, which is linked to checkpoint proteins"

### Supplemental Figure 1-6

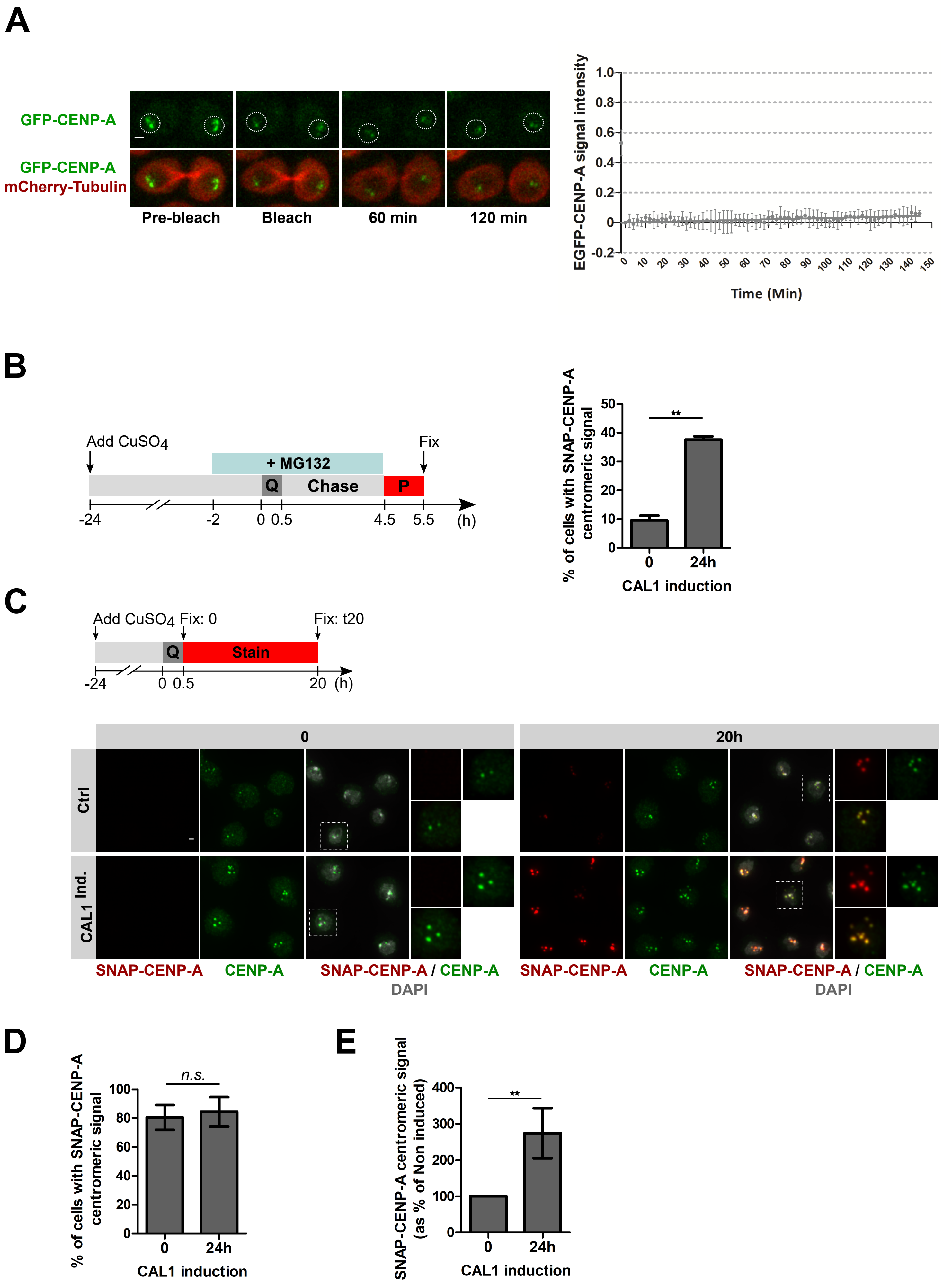

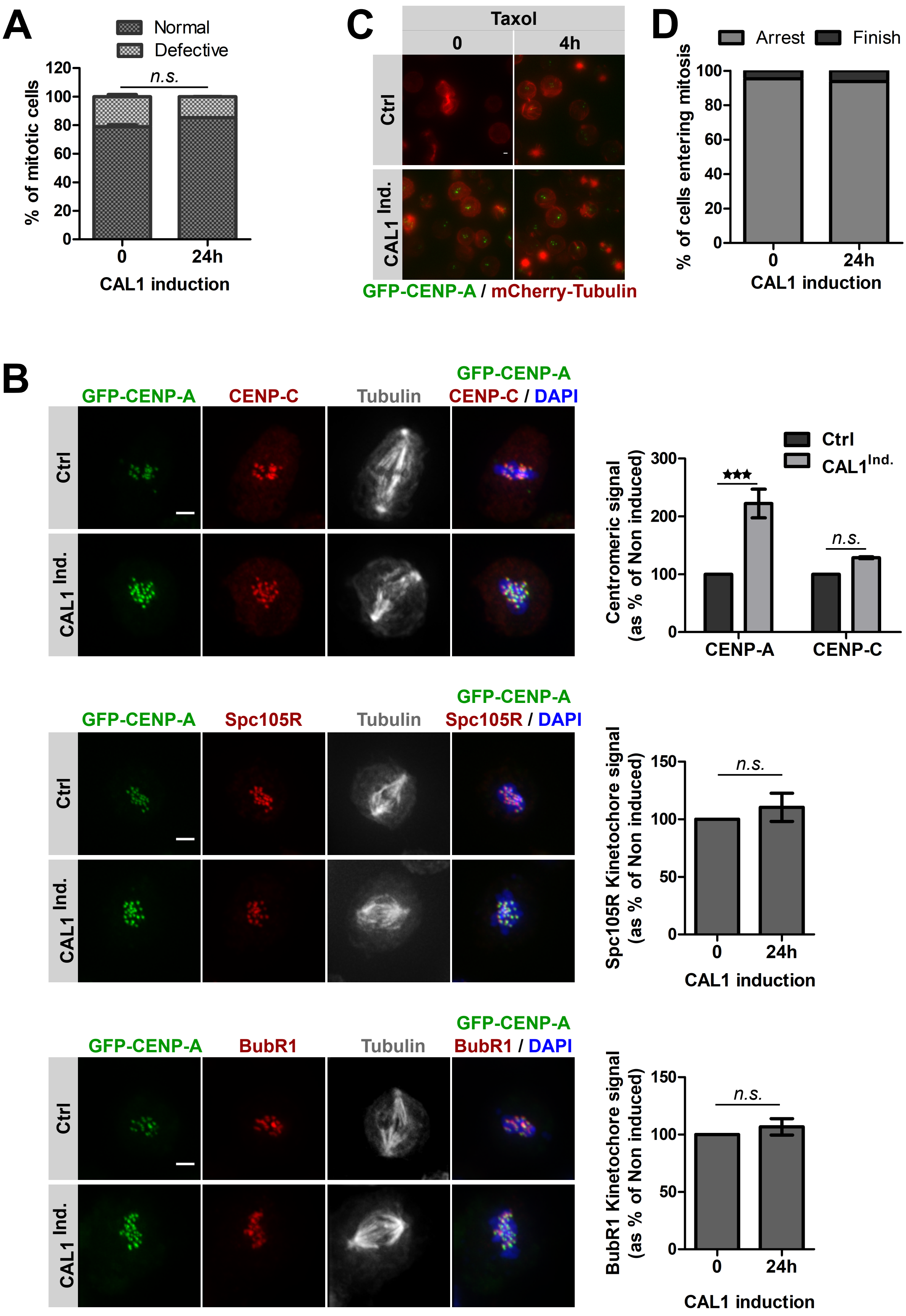

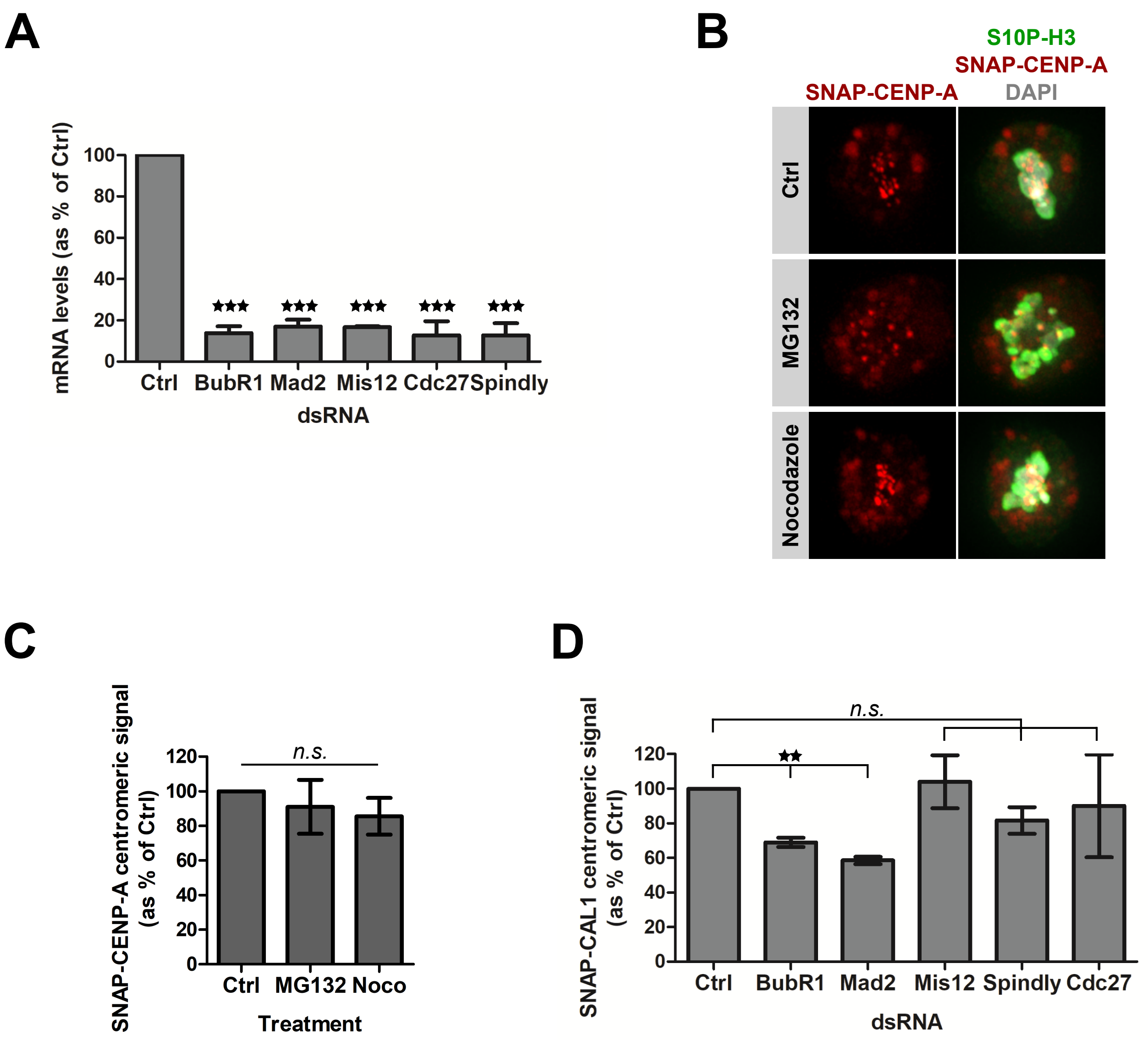

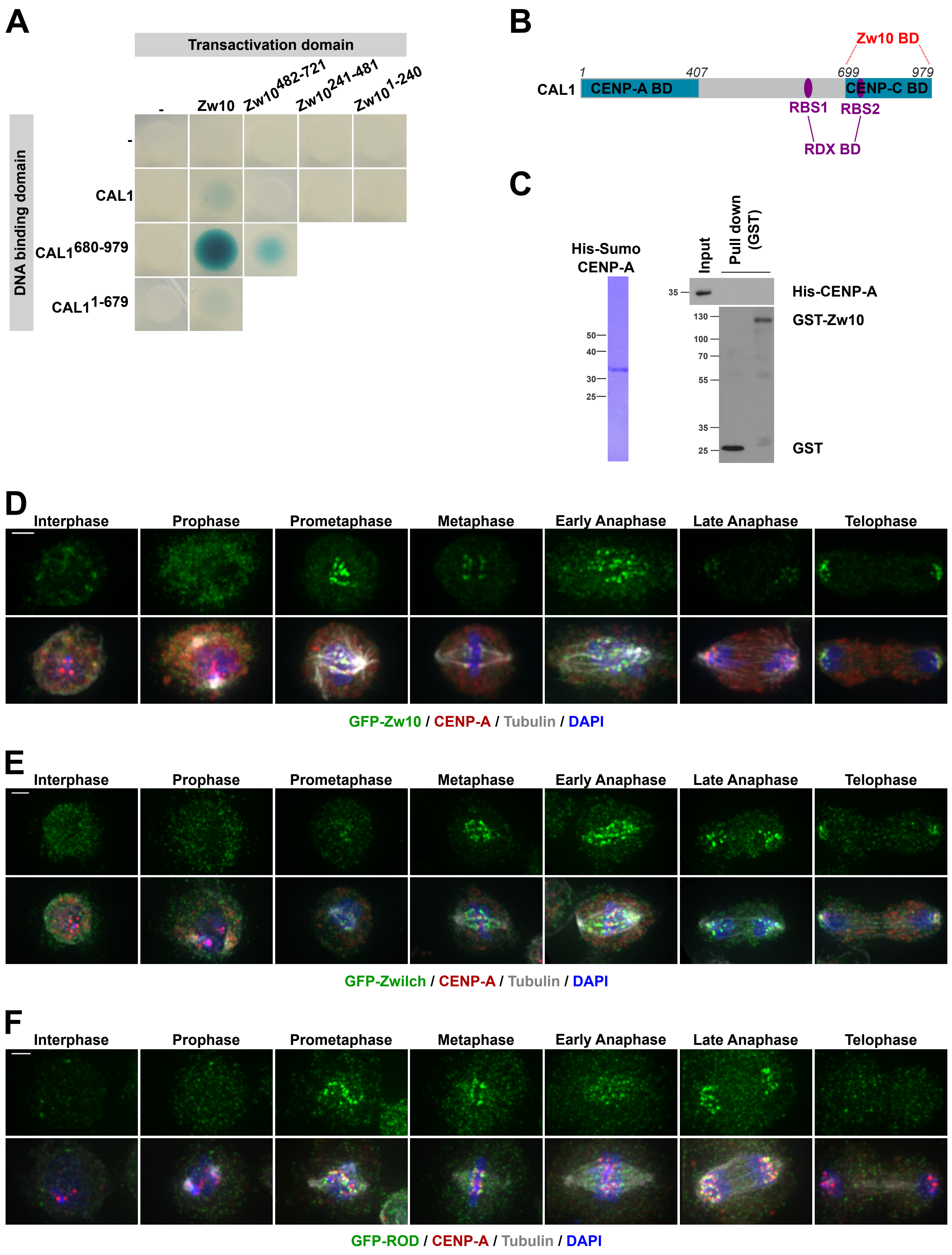

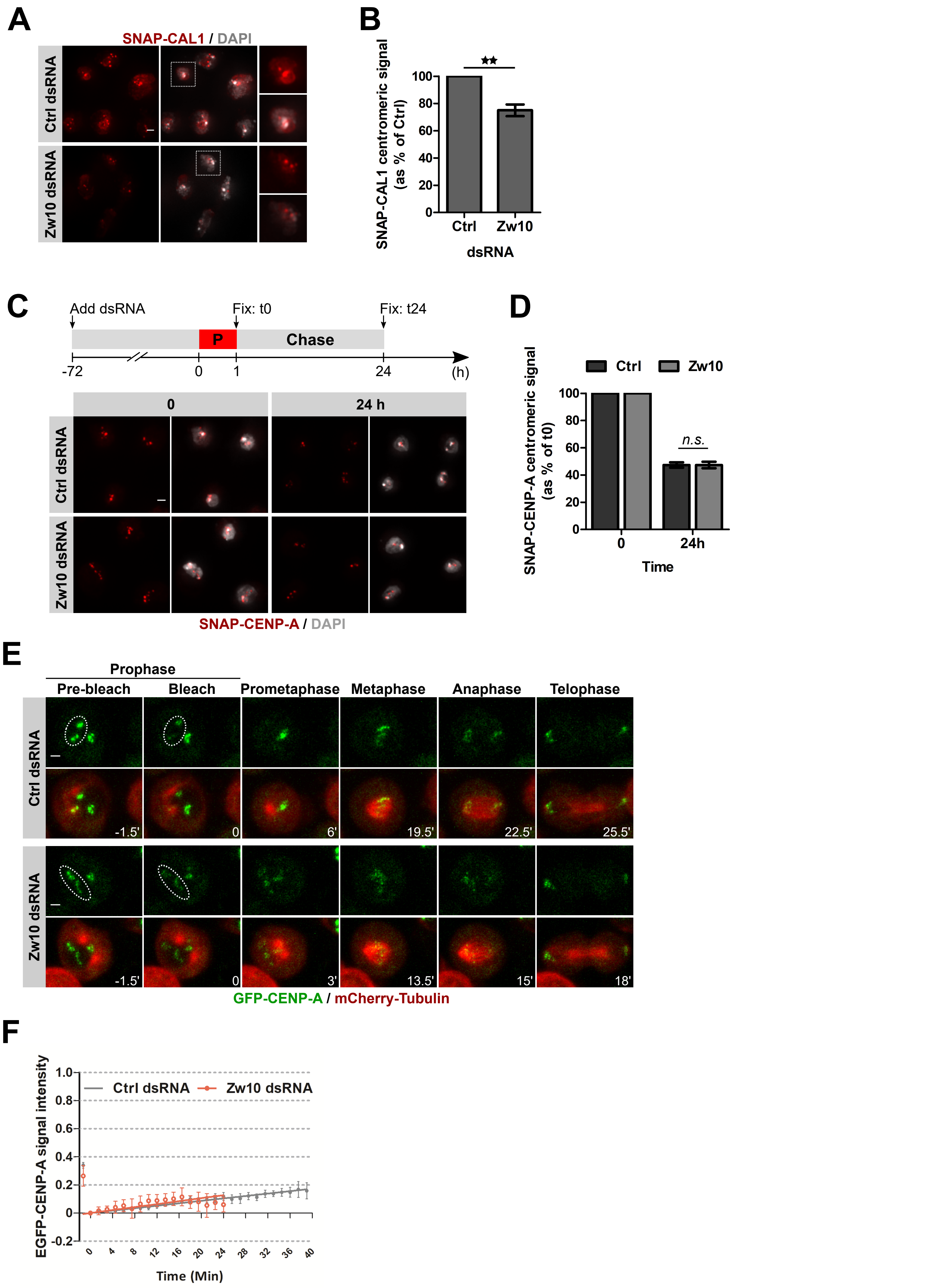

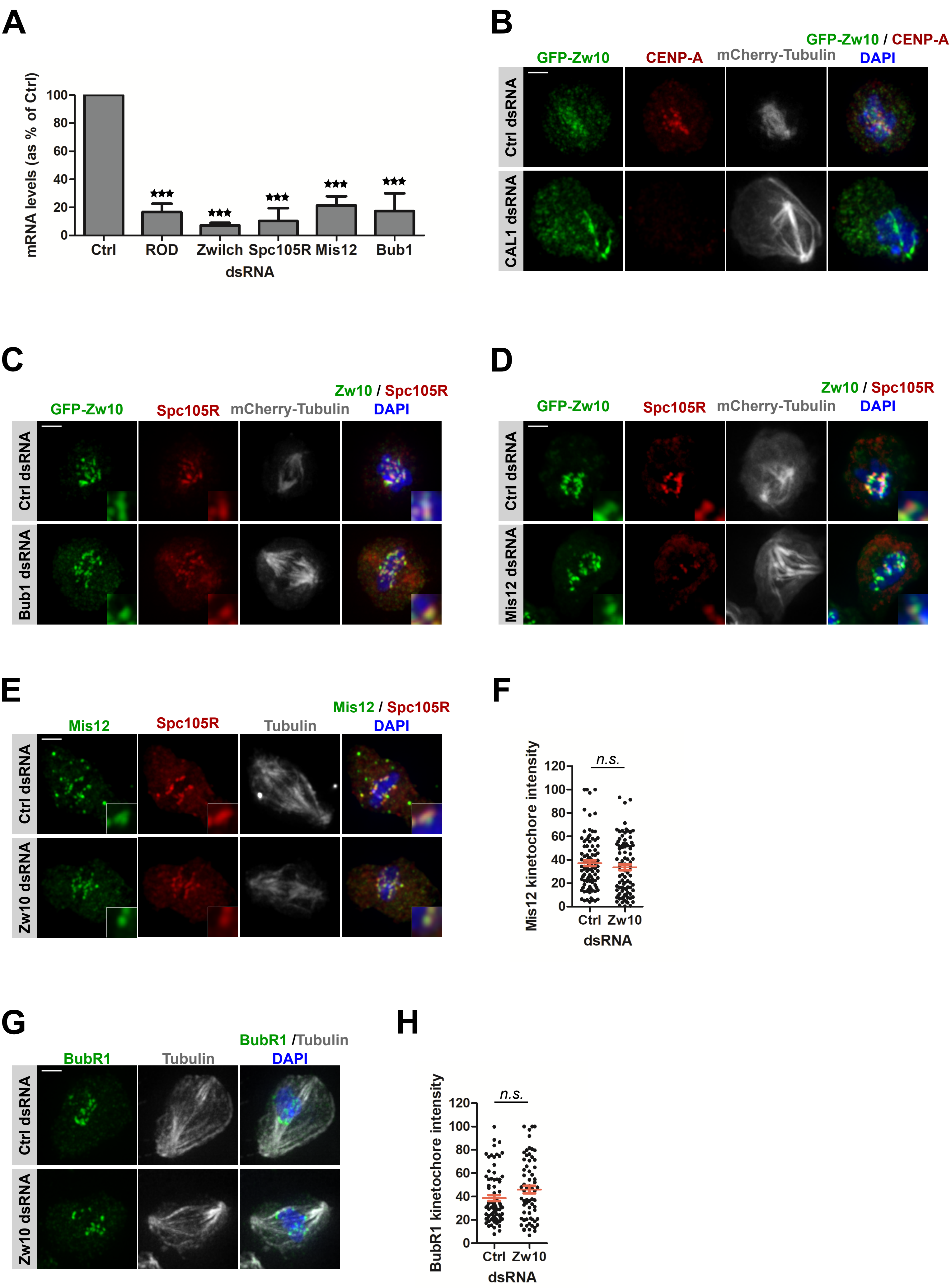
